# KHSRP-mediated Decay of Axonally Localized Prenyl-Cdc42 mRNA Slows Nerve Regeneration

**DOI:** 10.1101/2025.02.06.636857

**Authors:** M.D. Zdradzinski, Lauren S. Vaughn, Samaneh Matoo, Kayleigh Trumbull, Ashley Loomis, Elizabeth Thames, Seung Joon Lee, Nora Perrone-Bizzozero, Qun Lu, Jessica M. Larsen, J.L. Twiss

**Author notes:** **Corresponding Author:** Jeff Twiss Dept Biological Sciences 715 Sumter Street, CLS 401 University of South Carolina Columbia, SC 29204 USA, email –.

## Abstract

The small GTPase CDC42 promotes axon growth through actin filament polymerization and this growth is driven by axonal localization of the mRNA encoding the prenylated CDC42 isoform (*Prenyl-Cdc42*). Here, we show that axonal *Prenyl-Cdc42* mRNA transport and translation are decreased by growth-inhibiting stimulation and increased by growth-promoting stimulation. In contrast, axonal *RhoA* mRNA transport and translation are increased by growth inhibition but unaffected by growth promotion. Localized increase in KHSRP in response to growth inhibitory stimulation, through elevation of intracellular Ca^2+^, promotes decay of axonal *Prenyl-Cdc42* mRNA. Distinct 3’UTR motifs regulate transport and stability of axonal *Prenyl-Cdc42* mRNA. KHSRP protein binds to a *Prenyl-Cdc42* mRNA motif within nt 801-875 and the mRNA is remarkably increased in axons of *Khsrp^-/-^* mice. Selective depletion of *Prenyl-Cdc42* mRNA from axons reverses the accelerated axon regeneration seen in *Khsrp^-/-^*mice.

## INTRODUCTION

Polymerization and depolymerization of actin filaments in axonal growth cones, through the activation of small Rho GTPases, promotes axon extension, turning, and retraction (Luo et al., 1997). In a growing axon, intracellular signaling cascades from attractant and repulsant stimuli differentially activate Rho GTPases, including RHOA, RAC1, and CDC42 locally in growth cones. Activation of RHOA protein promotes actin filament depolymerization while activation of RAC1 and CDC42 proteins promotes actin filament polymerization (Hall, 1998). Synthesis of proteins locally in axons has emerged as a mechanism that can impact axon growth, both during development and after injury (Dalla Costa et al., 2020). Interestingly, the mRNAs encoding RHOA, RAC1, and a CDC42 isoform have been shown to localize into axons (Lee et al., 2021; Scott-Solomon and Kuruvilla, 2020; Wu et al., 2005). Translation of axonal *RhoA* mRNA in embryonic sensory neurons was initially reported in response to the growth inhibitory stimulus Semaphorin 3A, leading to growth cone collapse and axon retraction (Walker et al., 2012b; Wu et al., 2005). Conversely, nerve growth factor (NGF) activates axonal *Rac1* mRNA translation in developing sympathetic axons, with subsequent local prenylation of new RAC1 protein promoting axon extension (Scott-Solomon and Kuruvilla, 2020). The *CDC42* gene product is differentially spliced to generate prenylated- or palmitoylated-CDC42 proteins (Prenyl-CDC42 and Palm-CDC42, respectively) through mRNA isoforms that encode different C-termini and have different 3’ untranslated regions (UTR) We recently showed that the *Prenyl-Cdc42* mRNA, but not *Palm-Cdc42* mRNA, localizes into growing axons and is needed for axon growth (Lee et al., 2021). Taken together, these findings point to primary roles for intra-axonal synthesis of Rho GTPase proteins in axon extension and retraction but is not clear if and how the stoichiometry of the encoded proteins is determined by the balance of growth-promoting/attractant *vs.* growth-inhibiting/repulsive stimuli.

*Palm-Cdc42* and *Prenyl-Cdc42* mRNAs are generated by differential inclusion of *CDC42* gene’s exons 6 and 7 (Chen et al., 2012). Yap et al. (2016) reported that Palm-CDC42 (CDC42-Ex6) promotes dendritic spine development while Prenyl-CDC42 (CDC42-Ex7) promotes axon growth (Yap et al., 2016). Exons 6 and 7 encode for different C-terminal 10 amino acids in their protein products and the transcripts have different 3’UTR. We recently showed that the different 3’UTRs are responsible for distinct subcellular localization of these *Cdc42* mRNA isoforms in cortical neurons, with *Palm-Cdc42* mRNA localizing selectively into dendrites and *Prenyl-Cdc42* mRNA localizing into axons and dendrites (Lee et al., 2021). In dorsal root ganglion (DRG) sensory neurons that only extend axonal processes (Smith and Skene, 1997; Vuppalanchi et al., 2009; Zheng et al., 2001), *Prenyl-Cdc42* mRNA localizes into axons while *Palm-Cdc42* mRNA is retained in the cell body (Lee et al., 2021). In both sensory and cortical neurons, axon growth promotion by CDC42 was solely through the intra-axonally synthesized Prenyl-CDC42. Here, we show that the *Prenyl-Cdc42* mRNA 3’UTR contains two distinct motifs that impact axonal CDC42 levels. The proximal 35 nucleotides (nt) are needed for axonal localization, so this regulates how much of the mRNA localizes into axons. Just distal to that motif, an adenine-uridine (AU)-rich sequence modulates the levels of the mRNA within the axons, so that AU-rich-like sequence may functions as a stability motif. KHSRP protein binds to this AU-rich motif and promotes decay of axonal *Prenyl-Cdc42* mRNA, with axonal levels of *Prenyl-Cdc42* mRNA remarkably elevated in neurons lacking KHSRP. Furthermore, exposure to growth-inhibiting chondroitin sulfate proteoglycan (CSPG) increases axonal RHOA and KHSRP proteins and decreases axonal *Prenyl-Cdc42* mRNA’s transport and translation. Together, our findings show that axonal KHSRP drives decay of *Prenyl-Cdc42* mRNA, thereby slowing axon regeneration in the peripheral nervous system (PNS).

## RESULTS

### Extracellular stimuli can regulate axonal levels and translation of Prenyl-Cdc42 and RhoA mRNAs

We previously found that localized translation of *Prenyl-Cdc42* mRNA promotes neurite growth in cultured neurons (Lee et al., 2021). Since axon growth can be positively or negatively impacted by external stimuli, we asked if growth-promoting neurotrophins or a growth-inhibiting CSPG might alter axonal transport of *Prenyl-Cdc42* mRNA. For this, adult mouse DRG cultures were treated with either a neurotrophin cocktail, consisting of NT3, BDNF, and NGF to stimulate all 3 Trk receptors on DRG neuron subpopulations, or the CSPG aggrecan. smFISH for endogenous mRNAs showed that neurotrophin treatment increased and aggrecan decreased axonal levels of *Prenyl-Cdc42* mRNA (**Figure 1A-B; Suppl. Figure S1A**). In contrast to CDC42’s promotion of actin filament polymerization, RHOA activation causes actin filament depolymerization and the protein’s activity is increased by growth-inhibiting stimuli (Hall, 1998). *RhoA* mRNA also localizes into axons, and this was shown to increase upon exposure to CSPGs (Walker et al., 2012a). Consistent with Walker *et al*. (2012), we see that axonal *RhoA* mRNA levels increase in response to aggrecan; however, neurotrophin stimulation had no effect on axonal *RhoA* mRNA levels (**Figure 1A,C**). Together, these data point to reciprocal regulation of axonal *RhoA* and *Prenyl-Cdc42* mRNAs in response to CPSG stimulation but selective increase in axonal *Prenyl-Cdc42* mRNA in response to neurotrophins.

**Figure 1:**
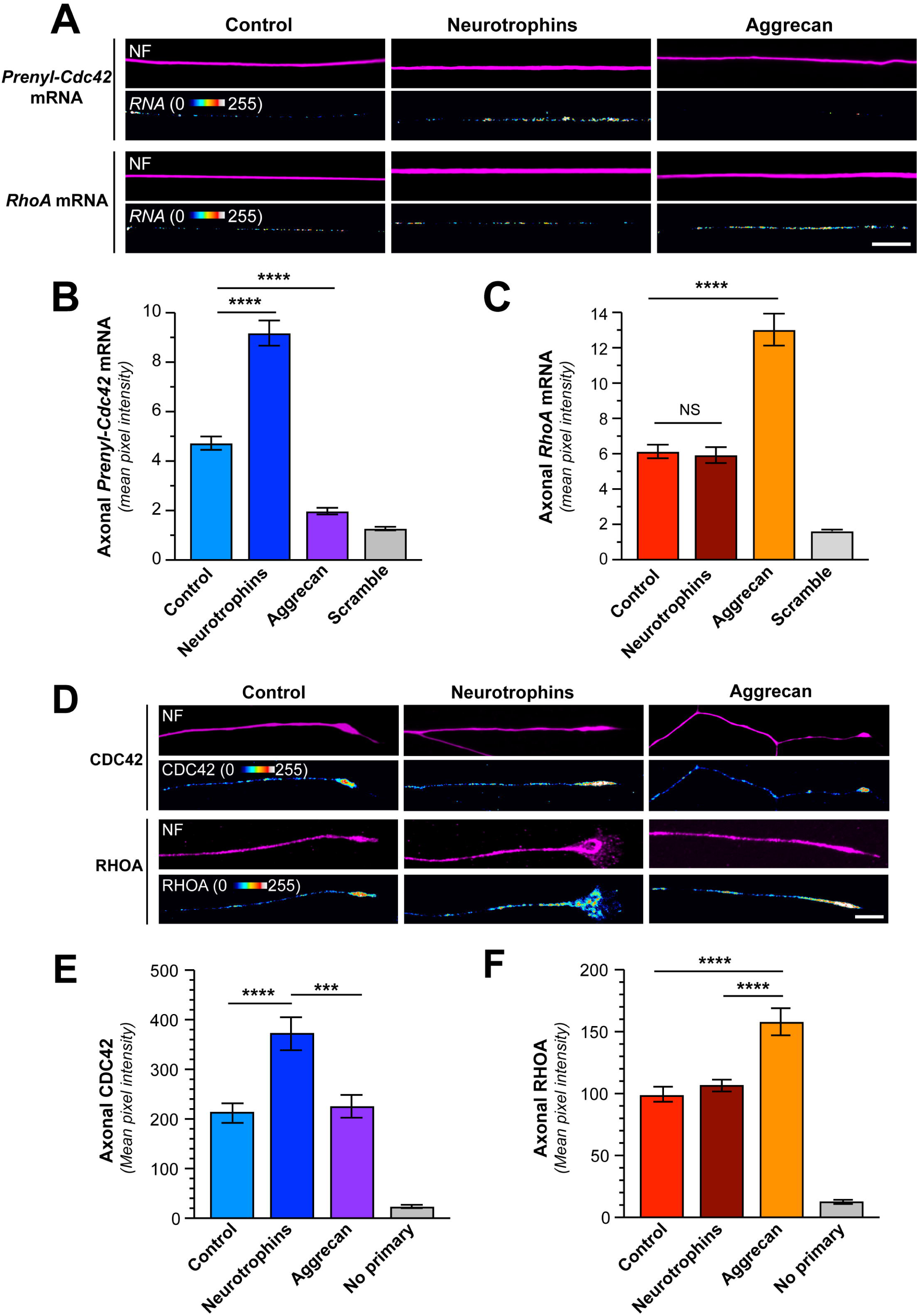
Growth-promoting and inhibiting stimuli regulate axonal levels of Prenyl-Cdc42 and RhoA mRNAs. **A)** Representative exposure-matched smFISH and IF images for *Prenyl-Cdc42* or *RhoA* mRNA respectively, and NF in adult DRG neuron cultures treated with either 10 ng/ml each NT3, BDNF, plus NGF each (‘neurotrophins’) or 50 ng/ml aggrecan [Scale bar = 10 µm] **B-C**) Quantification of smFISH signal intensities shown as mean pixel intensity above background for axons ± SEM (N ≥ 40 neurons across three independent cultures; *** P *<* 0.005, **** P < 0.001 by one-way ANOVA with pair-wise comparison with Tukey post-hoc tests). **D**) Representative exposure-matched IF images for Prenyl-CDC42 or RHOA proteins and NF in adult DRG neuron cultures treated with either 10 ng/ml neurotrophins or 50 ng/ml aggrecan; see Suppl. Figure S1A for representative no primary IF images [Scale bar =10 µm] **E-F**) Quantitation of axonal CDC42 (E) or RHOA (F) IF signal intensities shown as mean pixel intensity above background ± SEM; see Suppl, Figure S1B for cell body levels of CDC42 and RHOA under these conditions (N ≥ 40 neurons in three independent cultures; *** P < 0.005, **** P < 0.001 by one-way ANOVA with pair-wise comparison with Tukey post-hoc tests).

To determine if the endogenous Prenyl-CDC42 and RHOA proteins might show similar changes in their axonal levels with these stimuli, we performed immunofluorescence analyses on DRG cultures that had been treated with neurotrophins or aggrecan. Exposure to neurotrophins increased axonal CDC42 protein levels but had no apparent effect on axonal RHOA levels (**Figure 1D-F**). Aggrecan treatment increased RHOA protein levels in the axons but had no apparent effect on axonal CDC42 protein levels (**Figure 1D-E**). However, it should be noted that the CDC42 antibody utilized here does not distinguish between Prenyl-CDC42 and Palm-CDC42 proteins. We previously showed that Palm-CDC42 protein is transported into axons (Lee et al., 2021), so these immunofluorescence data do not distinguish between the two CDC42 isoforms. To more selectively assess axonal translation of *Prenyl-Cdc42* and *RhoA* mRNAs in response to these stimuli, we visualized axonal signals of diffusion limited GFP^MYR^ and mCherry^MYR^ reporters with the 5’ and 3’UTRs of rat *Prenyl-Cdc42* and *RhoA* mRNAs, respectively, as surrogates for local translation of the endogenous axonal mRNAs (GFP^MYR^5’/3’prenyl-Cdc42, mCherry^MYR^5’/3’RhoA). The 5’ and 3’UTRs were included to ensure that we captured both axonal localizing motifs and any translational control motifs; the MYR tag limits diffusion of the newly synthesized GFP and mCherry proteins in neurites such that fluorescence recovery after photobleaching (FRAP) can be used to visualize sites of reporter mRNA translation (Aakalu et al., 2001; Lee et al., 2021; Yudin et al., 2008). FRAP analyses in DRG neurons co-expressing *GFP^MYR^5’/3’prenyl-Cdc42* and *mCherry^MYR^5’/3’RhoA* mRNAs showed fluorescent recovery within 15 min post-bleach that was attenuated by pretreatment with the protein synthesis inhibitor anisomycin, consistent with intra-axonal translation of the reporter mRNAs (**Figure 2B-C; Suppl. Figure S1B-C**). To determine if neurotrophin or aggrecan stimulation might affect translation of the *GFP^MYR^5’/3’prenyl-Cdc42* and *mCherry^MYR^5’/3’RhoA* mRNAs, we bath applied the neurotrophin cocktail or aggrecan for 30 min before photobleaching. The neurotrophin stimulation increased *GFP^MYR^5’/3’prenyl-Cdc42* mRNA translation in distal axons but, consistent with the axonal mRNA analyses above, had no effect on axonal *mCherry^MYR^5’/3’RhoA* mRNA translation (**Figure 2A-C**). In contrast, aggrecan treatment decreased GFP^MYR^5’/3’prenyl-Cdc42 and increased mCherry^MYR^5’/3’RhoA fluorescence recovery (**Figure 2A-C**). The recovery of axonal GFP^MYR^5’/3’prenyl-Cdc42 fluorescence in the presence of aggrecan was largely indistinguishable from treatments with the protein synthesis inhibitor anisomycin (**Figure 2A-B; Suppl. Figure S1B**). Taken together, the data in Figures 1 and 2 are consistent with reciprocal regulation of axonal *Prenyl-Cdc42* and *RhoA* mRNA levels and subsequent intra-axonal translation in response to aggrecan. In contrast, responsiveness to neurotrophins appears to be limited to the axonal *Prenyl-Cdc42* mRNA.

**Figure 2:**
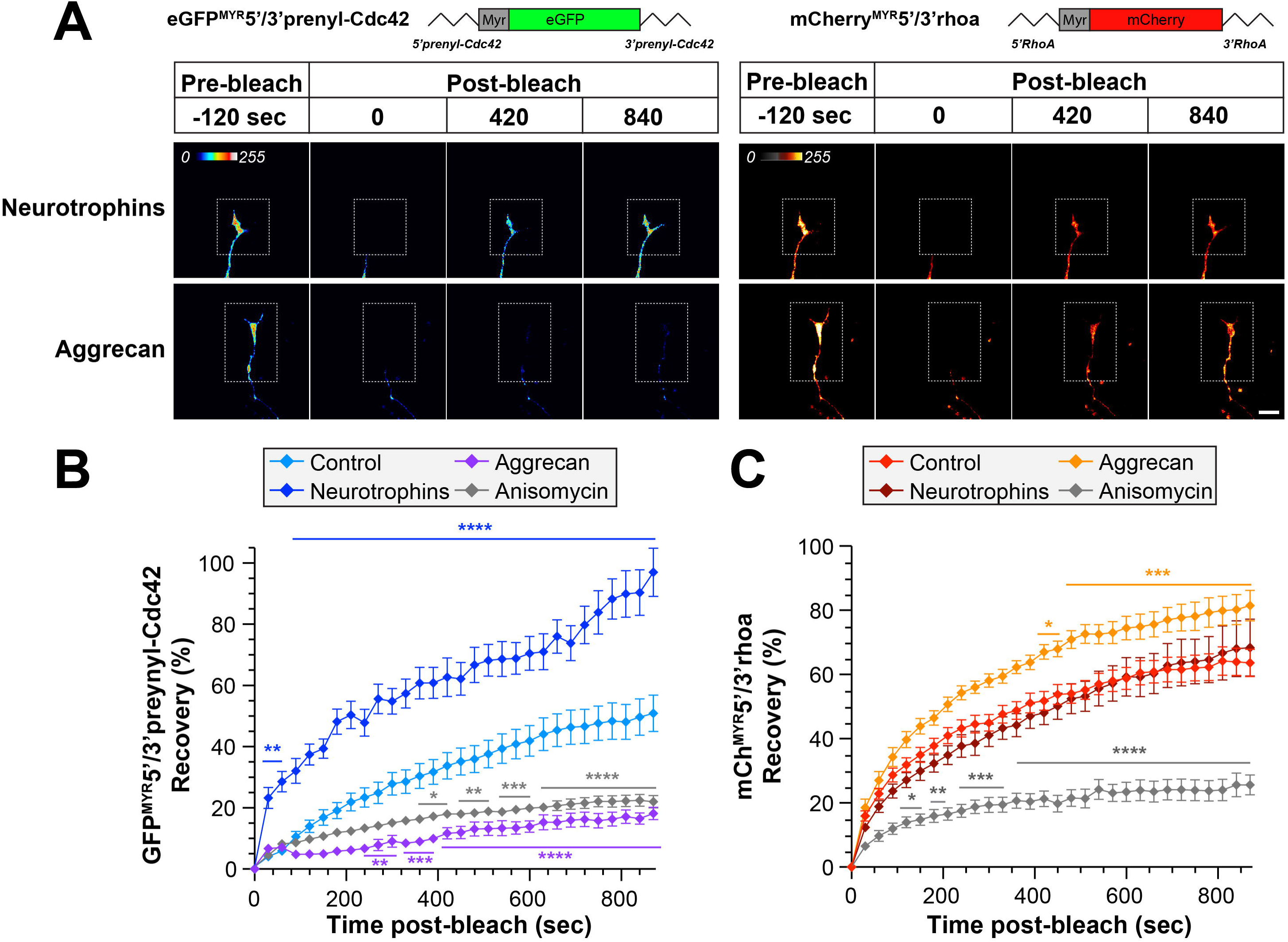
Growth-promoting and inhibiting stimuli regulate axonal translation of Prenyl-Cdc42 and RhoA mRNAs. **A)** Representative FRAP image sequences for DRG neurons co-transfected with GFP^MYR^5’/3’prenyl-cdc42, and mCherry^MYR^5’/3’rhoA (72 h post-transfection) are shown. Cultures were treated with either 10 ng/ml neurotrophins or 50 ng/ml aggrecan as in Figure 1. Boxed regions represent the photobleached ROIs; see Suppl. Figure S1C-D for control and anisomycin-treated representative FRAP image sequences [Scale bar = 20 µm]. **B-C**) Quantitation of FRAP sequences from panel A are shown as average normalized % recovery11±11 SEM. Note that translation inhibition with anisomycin prior to photobleaching shows that the *GFP^MYR^* and mCherry^MYR^ recovery requires protein synthesis (N ≥ 10 neurons over three independent experiments; * P < 0.05, \**\** P < 0.01, ***P < 0.005, ****P < 0.001 by two-way ANOVA with pair-wise comparison and Tukey post-hoc tests).

### An evolutionarily conserved region of the prenyl-Cdc42 mRNA 3’UTR drives its axonal localization

As noted above, *Prenyl-Cdc42* mRNA localizes into axons of sensory neurons and cortical neurons and the 3’UTR encoded by *CDC42* exon 7 is necessary and sufficient for the mRNA’s axonal localization (Lee et al., 2021). We have previously shown that conservation of 3’UTR regions can have predictive value for identifying functional domains in mRNAs (Lee et al., 2018; Vuppalanchi et al., 2010). The initial 150 nt of rat *Prenyl-Cdc42* 3’UTR (nt 764-913; NCBI XM_008764286.3) shows > 85 % sequence identity with the 3’UTRs of 26 other vertebrate species (**Figure S2**). To determine if this conserved 150 nt region contains a functional motif, we generated fluorescent reporter constructs containing *Prenyl-Cdc42*’s nt 764-913 or 914-2164 (*i.e.*, the remainder of the 3’UTR) as the 3’UTR for a myristoylated (MYR) GFP cDNA (GFP^MYR^3’prenyl-Cdc42^764-913^ and GFP^MYR^3’prenyl-Cdc42^914-2164^, respectively; **Figure 3A**). smFISH analyses of transfected adult mouse DRG cultures showed that *GFP^MYR^* mRNA only localized into axons of the GFP^MYR^3’prenyl-Cdc42^764-913^ transfected neurons; GFP^MYR^3’prenyl-Cdc42^914-2164^ transfected neurons did not show axonal *GFP* mRNA signal above the scrambled control probe (**Figure 3B,D; Suppl. Figure S3A**). Cell body levels of *GFP* mRNA were not appreciably different between GFP^MYR^3’prenyl-Cdc42^764-913^ and GFP^MYR^3’prenyl-Cdc42^914-2164^ transfected neurons (**Figure 3B; Suppl. Figure S3B**). These data show that the conserved proximal 150 nt region of *Prenyl-Cdc42*’s 3’UTR is necessary and sufficient to drive axonal mRNA localization.

**Figure 3:**
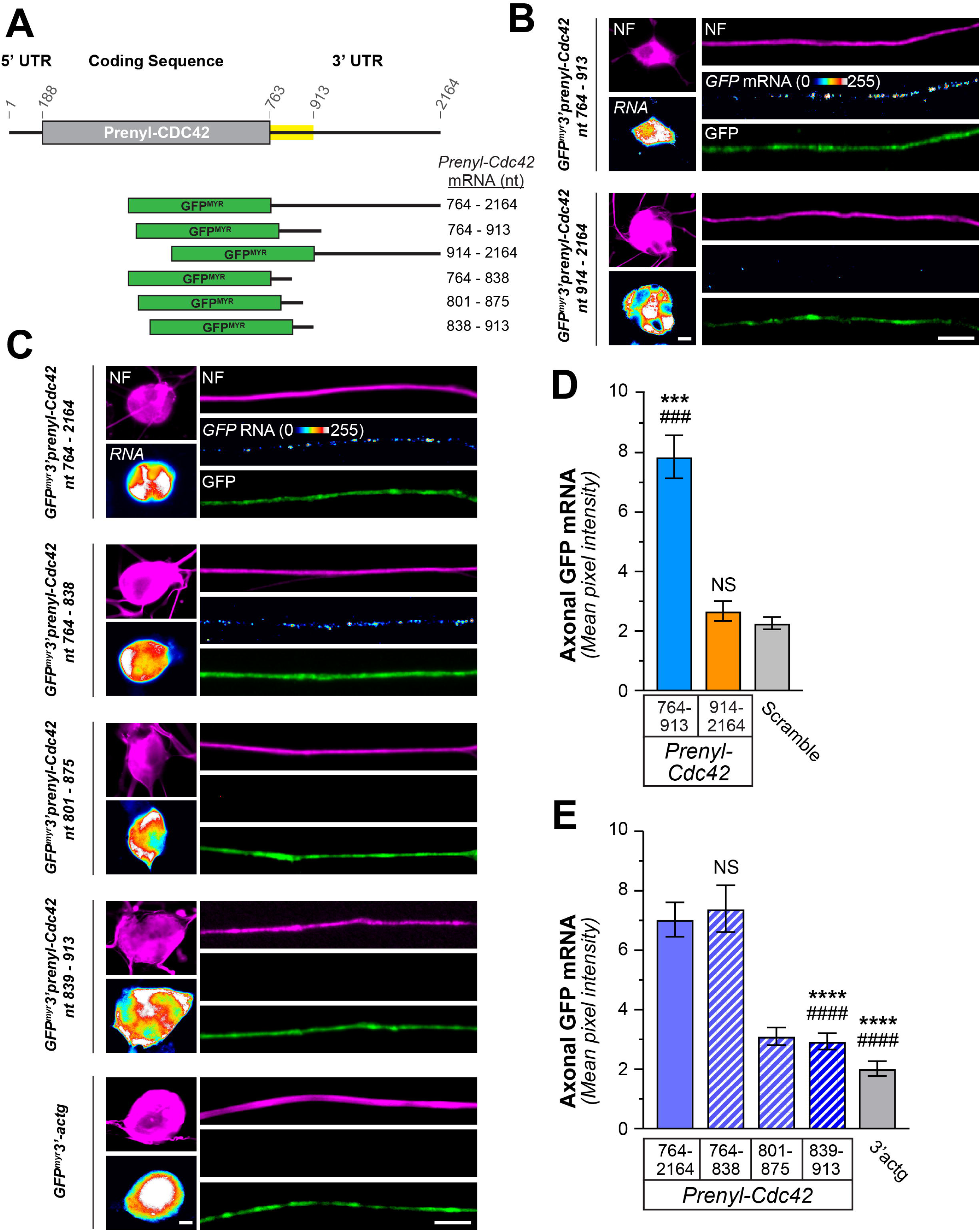
An evolutionarily conserved region of the Prenyl-Cdc42 mRNA 3’ UTR drives its axonal localization. **A)** Schematic of the regions of the rat *prenyl*-*Cdc42* 311UTR mRNA that were tested for axonal localizing activity. The yellow-shaded regions show ≥85% sequence identity between available mammalian *prenyl*-*Cdc42* mRNAs (see Suppl. Figure S2 for sequence alignments). **B)** Representative exposure-matched smFISH and immunofluorescence (IF) images for GFP^myr^ mRNA and neurofilament (NF) in adult DRG neuron cultures transfected with GFP^MYR^3’prenyl-Cdc42^764-913^, and GFP^MYR^3’prenyl-Cdc42^914-2164^. See Suppl. Figure S3A for representative images of scrambled smFISH probe [Scale bar = 10 µm]. **C)** Representative exposure-matched smFISH and IF images for GFP mRNA plus NF in adult DRG neuron cultures transfected with GFP^MYR^3’prenyl-Cdc42^764-2164^, GFP^MYR^3’prenyl-Cdc42^764-838^, GFP^MYR^3’prenyl-Cdc42^801-875^, GFP^MYR^3’prenyl-Cdc42^839-913^, or GFP^MYR^3’actg [Scale bar = 10 µm] **D-E**) Quantitation of smFISH signal intensities shown as mean ± standard error of the mean (SEM) pixel intensity above background for axons. Corresponding cell body mRNA levels are shown in Suppl. Figure S3B-C (N ≥ 40 neurons across three independent cultures; *** P < 0.005 compared to 913-2164 and ### P < 0.005 compared to scramble in D and **** P < 0.001 compared to 764-838 and #### P < 0.001 compared to 764-2164 in E by one-way ANOVA with pair-wise comparison and Tukey post-hoc tests).

801-875 nt of the *Prenyl-Cdc42* mRNA contains approximately 80% adenine and uridine bases, which could represent an AU-rich element (ARE) (Barreau et al., 2005). KHSRP and ELAV-like protein 4 (HuD), which both localize into CNS and PNS axons, have been shown to bind to axonal ARE-containing mRNAs and both proteins have been implicated in subcellular mRNA localization (Akten et al., 2011; Rehbein et al., 2000; Rehbein et al., 2002; Yoo et al., 2013). The ARE in *Gap43* mRNA’s 3’UTR is necessary and sufficient for its localization into rat sensory axons (Yoo et al., 2013). Thus, we asked if the AU rich region in *Prenyl-Cdc42* nt 764-913 is sufficient for axonal localization of the mRNA. For this, we generated GFP^MYR^ reporter constructs containing 3 overlapping 75 nt portions of the 764-913 sequence (GFP^MYR^3’prenyl-Cdc42^764-838^, GFP^MYR^3’prenyl-Cdc42^801-875^, and GFP^MYR^3’prenyl-Cdc42^839-913^; **Figure 3A**). The 3’UTR of *GFP^MYR^3’prenyl-Cdc42^801-875^* mRNA is quite AU-rich while the *GFP^MYR^3’prenyl-Cdc42^764-838^* contains a short segment of the AU-rich region only at its 3’ end. DRGs expressing the *GFP^MYR^3’prenyl-Cdc42^764-838^* mRNA showed axonal localization of *GFP* mRNA by smFISH, but axonal *GFP* mRNA signals in the *GFP^MYR^3’prenyl-Cdc42^801-875^* and *GFP^MYR^3’prenyl-Cdc42^839-913^*expressing neurons were not distinguishable from neurons transfected with GFP^MYR^ containing a non-localizing 3’UTR (GFP^MYR^3’Actg; **Figure 3C,E**). Cell body levels of *GFP* mRNA were not significantly different between GFP^MYR^3’prenyl-Cdc42^764-838^, GFP^MYR^3’prenyl-Cdc42^801-875^, GFP^MYR^3’prenyl-Cdc42^839-913^, and GFP^MYR^3’actg transfected neurons (**Figure 3C, Suppl. Figure S3C**). Taken together, these data indicate that nt 764-838 of *Prenyl-Cdc42* mRNA, the proximal 75 nt of its 3’UTR, is necessary and sufficient for the mRNA’s transport into axons.

### ARE-binding protein KHSRP binds to non-localizing conserved region of Prenyl-Cdc42 3’UTR

The decrease in axonal *Prenyl-Cdc42* mRNA in response to aggrecan coupled with the presence of an AU-rich region in its 3’UTR raise the possibility that ARE binding proteins could impact *Prenyl-Cdc42* mRNA levels in the axons. KHSRP binds ARE-containing mRNAs to promote their decay by targeting them to the cytoplasmic exosome (Gherzi et al., 2004). Consistent with this, *Gap43* mRNA is stabilized in PNS axons of KHSRP knockout mice (Patel et al., 2022). Thus, we tested whether KHSRP might bind to the 3’UTR of *Prenyl-Cdc42* mRNA. We initially asked whether *Prenyl-Cdc42* mRNA forms a complex with KHSRP using RNA co-immunoprecipitation (RIP). Anti-KHSRP antibody immunoprecipitated KHSRP from DRG culture lysates but KHSRP was not detected in immunoprecipitations performed with a non-reactive rabbit IgG (**Figure 4A**). RNA isolated from these immunoprecipitates showed that approximately 25% of input *Prenyl-Cdc42* mRNA co-precipitated with KHSRP by reverse transcriptase-coupled droplet digital PCR (RTddPCR; **Figure 4B**). Thus, endogenous KHSRP and *Prenyl-Cdc42* mRNA exists in a complex within PNS dorsal root ganglion.

**Figure 4:**
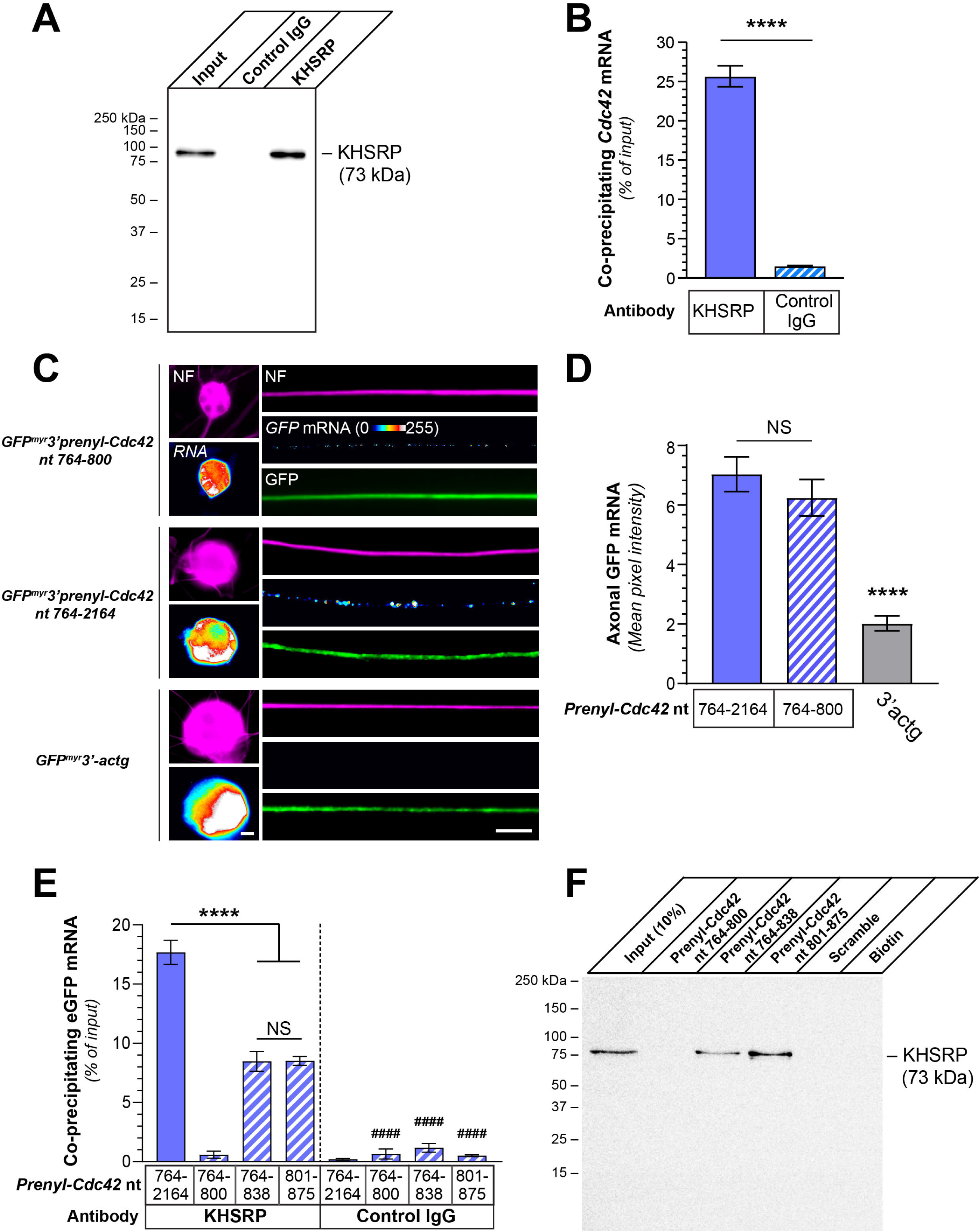
ARE-binding protein KHSRP binds to non-localizing conserved region of Prenyl-Cdc42 3’UTR. **A-B**) Representative western blot for KHSRP protein using KHSRP *vs*. control IgG immunoprecipitation from adult DRG cultures is shown in A. B shows RTddPCR analyses of RNA isolated from KHSRP vs. control IgG immunoprecipitates. Co-precipitating *Prenyl-Cdc42* mRNA is shown as mean mRNA copies as percentage of input ± SEM (N = 3 biological replicates; **** P ≤ 0.001 by Student’s t-test for the indicated data pairs). **C)** Representative exposure-matched smFISH and IF images for *GFP* mRNA and NF in adult DRG neuron cultures transfected with GFP^MYR^3’prenyl-Cdc42^764-2164^, GFP^MYR^3’prenyl-Cdc42^764-800^, or GFP^MYR^3’actg [Scale bar = 10 µm]. **D)** Quantitation of smFISH signal intensities shown as mean pixel intensity above background for axons ± SEM; see Suppl. Figure S4A for cell body RNA levels (N ≥ 40 neurons in three independent cultures; NS = not significant, **** P < 0.001 compared to 764-2164 (one-way ANOVA with pair-wise comparison with Tukey post-hoc tests). **E)** Analyses of *GFP* mRNA from KHSRP and control IgG immunoprecipitates from DRG neuron cultures transfected with GFP^MYR^3’prenyl-Cdc42^764-2164^, GFP^MYR^3’prenyl-Cdc42^764-800^, GFP^MYR^3’prenyl-Cdc42^764-838^, or GFP^MYR^3’prenyl-Cdc42^801-875^ is shown as mean of co-precipitating mRNA copies as percentage of input ± SEM (N = 3 biological replicates; **** P ≤0.001 for indicated data pairs and #### P ≤ 0.001 for control IgG compared to corresponding KHSRP RIP by two-way ANOVA with pair-wise comparison and Tukey post-hoc tests). **F)** Representative immunoblot analysis for KHSRP protein in RNA affinity pull down using biotinylated oligonucleotides corresponding to nt 764-838, 801-875 and 764-800 of rat *Prenyl-Cdc42* mRNA. Scrambled oligonucleotide and biotin alone were used as negative controls. See Suppl. Figure S5 for analysis of *Prenyl-Cdc42* 3’UTR RNA oligonucleotides binding to recombinant KHSRP protein *in vitro* (N = 3 independent pull downs).

Since nt 764-838 in the *Prenyl-Cdc42* mRNA sequence is sufficient for the mRNA’s localization into axons (see Figure 3C,E) and *Prenyl-Cdc42*’s AU-rich sequence begins at nt 801, we asked whether the RNA region forming a complex with KHSRP might be the same as that needed for axonal localization of *Prenyl-Cdc42* mRNA. For this, we generated a fluorescent reporter construct without the AU-rich region in the *Prenyl-Cdc42*’s 3’UTR (GFP^MYR^3’prenyl-Cdc42^764-800^) to distinguish potential functions nt 764-800 and 801-875. DRG neurons expressing *GFP^MYR^3’Prenyl-Cdc42^764-800^* mRNA showed robust axonal *GFP* FISH signals, which were comparable to values seen with the *GFP^MYR^3’prenyl-Cdc42^764-838^* mRNA expressing neurons shown in Figure 1 (**Figure 4C-D**). As expected, the GFP^MYR^3’actg transfected neurons showed no detectable *GFP* mRNA and analyses of cell bodies showed comparable expression of *GFP^MYR^3’prenyl-Cdc42^764-800^* and *GFP^MYR^3’actg* mRNAs (**Figure 4C, Suppl. Figure S4A**). Thus, the axonal localization motif in *Prenyl-Cdc42* mRNA lies in the most proximal 37 nt of its 3’UTR (nt 764-800) and is functionally distinct from the more 3’ AU-rich sequence that begins at nt 801.

We next sought to determine if *Prenyl-Cdc42* mRNA nt 801-875 comprises a functional motif by asking if KHSRP interacts with the RNA sequence. For this, we performed KHSRP RIP analyses from DRG cultures transfected with the GFP^MYR^3’prenyl-Cdc42^764-2164^, GFP^MYR^3’prenyl-Cdc42^764-800^, GFP^MYR^3’prenyl-Cdc42^764-838^, or GFP^MYR^3’prenyl-Cdc42^801-875^ expression constructs. RTddPCR analyses of the immunoprecipitates showed that *GFP^MYR^* mRNAs containing *Prenyl-Cdc42* mRNA nt 764-2164, 764-838, and 801-875, but not 764-800, were precipitated by anti-KHSRP antibodies (**Figure 4E**). IgG control immunoprecipitation showed no detectable *GFP^MYR^* mRNA for any of the DRG transfectants (**Figure 4E**). The levels of *GFP^MYR^* mRNA coprecipitating with KHSRP in the *GFP^MYR^3’prenyl-Cdc42^764-838^* and *GFP^MYR^3’prenyl-Cdc42^801-875^* mRNA expressing DRG cultures were about half of what was seen in the *GFP^MYR^3’prenyl-Cdc42^764-2164^* mRNA expressing cultures suggesting that KHSRP interaction may be affected by sequences downstream of nt 875 (**Figure 4E**). Consistent with this, ARE-like sequences are present downstream of nt 864 in the *Prenyl-Cdc42* mRNA 3’UTR (*e.g.,* nt 994-1003). We used an *in vitro* RNA affinity pulldown technique where biotinylated RNA oligonucleotides serve as ‘bait’ to test whether endogenous proteins from sciatic nerve axoplasm isolates can bind to the RNA bait (Doron-Mandel et al., 2016). Endogenous KHSRP was clearly detected by immunoblotting in affinity pulldowns with the *Prenyl-Cdc42* mRNA nt 764-838 and 801-875 oligonucleotides but not with the nt 764-800 or scrambled oligonucleotide affinity pulldowns (**Figure 4F**). Together, these data show that KHSRP protein binds directly to the 3’UTR of *Prenyl-Cdc42* mRNA, which could account for the rapid decline of axonal *Prenyl-Cdc42* mRNA in response to aggrecan exposure.

### KHSRP deletion increases axonal Prenyl-Cdc42 mRNA

Axonal KHSRP levels increase in rodent sciatic nerve axons after crush injury via localized translation of *Khsrp* mRNA (Patel et al., 2022). Since CDC42 protein activity has been shown to increase axon growth (Lee et al., 2021; Luo et al., 1997) and the injury-induced increase in axonal KHSRP levels slows regeneration of peripheral nerves (Patel et al., 2022), we asked whether KHSRP might post-transcriptionally regulate *Prenyl-Cdc42* mRNA within peripheral nerve axons. For this, we compared *Prenyl-Cdc42* mRNA levels in axons of *Khsrp^-/-^* and *Khsrp^+/+^* mice (Lin et al., 2011) *in vivo* before and 7 days after a sciatic nerve crush injury. In contrast to our previous work where we did not detect *Prenyl-Cdc42* mRNA in uninjured, non-growing axons of wild type neurons (Lee et al., 2021), *Prenyl-Cdc42* mRNA was easily detected in the axons of uninjured *Khsrp^-/-^* sciatic nerves (**Figure 5A-B; Suppl. Figure S4B**). As anticipated, crush injury increased axonal *Prenyl-Cdc42* mRNA levels in the *Khsrp^+/+^* sciatic nerves (**Figure 5A-B**). Remarkably, the regenerating sciatic nerves of the *Khsrp^-/-^* mice showed approximately 5-fold higher axonal *Prenyl-Cdc42* mRNA signals compared to those of the *Khsrp^+/+^* mice (**Figure 5B**). Thus, KHSRP likely promotes decay of *Prenyl-Cdc42* mRNA in PNS axons under both naïve and regenerating conditions.

**Figure 5:**
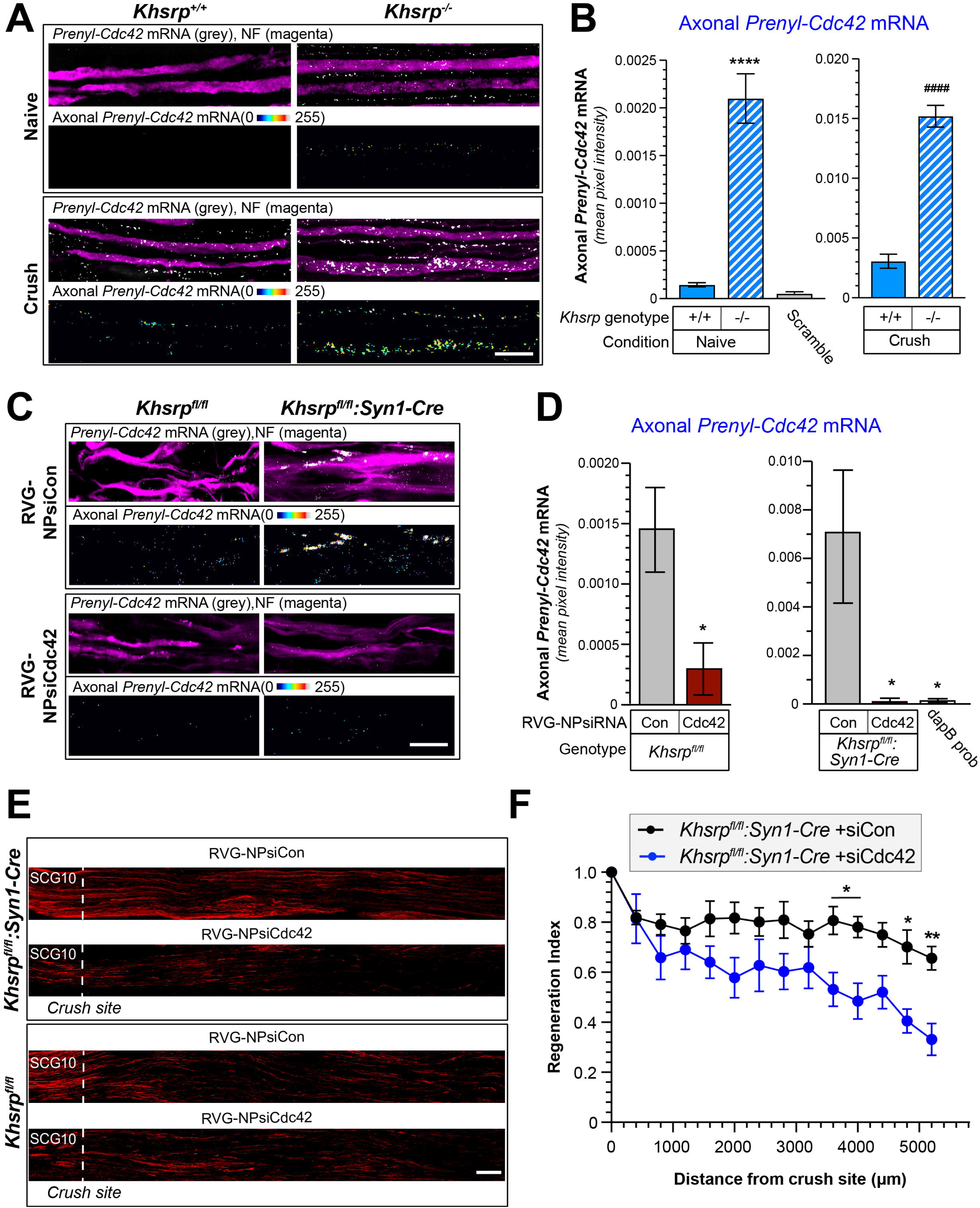
Prenyl-Cdc42 mRNA is increased in KHSRP knockout axons. **A)** Representative smFISH and IF images for naïve and 7 day post-crush injured sciatic nerve from *Khsrp^+/+^* or *Khsrp^-/-^* mice. The upper row of each image set shows the merged confocal *XY* optical plane; the lower row of each image set shows RNA signal overlapping with NF across individual optical planes that was extracted to a separate channel and projected as *XYZ* images; see Suppl. Figure S4B for representative images of scrambled smFISH probe [Scale bar = 5 µm]. **B)** Quantitation of smFISH signals for RNA probe signals overlapping with NF shown in A as mean ± SEM (N = 3 biological replicates; **** P < 0.001 for indicated data pairs by two-way ANOVA with pair-wise comparison and Tukey post-hoc tests) [Scale bar = 10 µm] **C-D)** Representative exposure-matched smFISH/IF for *Cdc42* mRNA in sciatic nerve axons of *Khsrp^fl/fl^:Syn1-Cre* vs. *Khsrp^fl/fl^* mice at 14 d post-crush shown in C. RVG-NPsiRNAs were applied at 7 d post-crush. Merged images show single XY planes and ‘axon only’ images show XYZ project of extracted RNA pixels that overlap with NF in individual Z planes. Quantification of axonal *Prenyl-Cdc42* mRNA levels shown in D; note that Y-axis scale is different for *Khsrp^fl/fl^* and *Khsrp^fl/fl^:Syn1-Cre* mice (N = 5 animals per condition; * P < 0.05 for indicated data pairs, as determined by an unpaired t-test) [Scale bar = 5 µm] **E-F)** Representative images for SCG10 immunostaining (E) and regeneration indices (F) for *Khsrp^fl/fl^:Syn1-Cre* mice treated with RVG-NPsiRNAs targeting *Prenyl-Cdc42* mRNA vs. non-targeting siRNA control are shown. See Suppl. Figure S4 D,E for representative images and quantitation of regeneration indices in *Khsrp^fl/fl^* mice treated with RVG-NPsiRNAs (N = 5 animals per condition; * P < 0.05 by repeated measures ANOVA) [Scale bar = 100 µm].

To test for potential functional significance of the elevated *Prenyl-Cdc42* mRNA in *Khsrp^-/-^* axons, we used an *in vivo* siRNA approach to selectively deplete *Prenyl-Cdc42* mRNA from sciatic nerve axons. We reasoned that delivering an siRNA directly to the sciatic nerve would allow us to selectively deplete *Prenyl-Cdc42* mRNA from sciatic nerve axons. Thus, we packaged siRNAs targeting *Prenyl-Cdc42* mRNA or non-targeting siRNA (siPrenyl-Cdc42 and siCon, respectively) into a polymersome nanoparticle (Trumbull et al., 2024); the exterior surface of the nanoparticles was tagged with 29 amino acid rabies virus glycoprotein peptide-9R (RVG) that has previously been used to deliver nanoparticle cargos to neurons (Kwon et al., 2016) and has been shown to bind to NCAM (Thoulouze et al., 1998). We confirmed neuronal uptake of the siRNA laden RVG-nanoparticles (RVG-NPsiRNA) in mouse DRG cultures (data not shown). The NP-siRNA were next tested for specificity *in vivo* by direct injection into the sciatic nerve of wild type mice that had undergone sciatic nerve crush 7 days previously to increase axonal *Prenyl-Cdc42* mRNA. Mice injected with RVG-NPsiRNA targeting *Prenyl-Cdc42* compared to non-targeting siRNA (siCdc42 and siCon, respectively) showed depletion of *Prenyl-Cdc42* mRNA based on RTddPCR analyses of sciatic nerve axoplasm (**Suppl. Figure S4C**). We next asked whether *Prenyl-Cdc42* mRNA depletion from axons of KHSRP-deficient mice could decrease the accelerated regeneration seen in mice lacking neuronal KHSRP. For this, we performed a sciatic nerve crush injury in *Khsrp^fl/fl^* crossed to *Syn1-Cre* (*KHSRP^fl/fl^:Syn1-Cre*) and *KHSRP^fl/fl^* mice (Olguin et al., 2022); 7 days later the sciatic nerve proximal to the crush site as well as the approximately same level in the contralateral (sham) nerve was injected with RVG-NPsiRNAs for *Prenyl-Cdc42* vs. non-targeting control. Both RT-ddPCR using RNA isolated from axoplasm and smFISH showed a significant depletion of axonal *Prenyl-Cdc42* mRNA in the siCdc42 vs. siCon polymersome-injected mice (**Figure 5C-D**). Nerve regeneration was significantly reduced by *Prenyl-Cdc42* mRNA depletion from the axons of the *Khsrp^fl/fl^:Syn1-Cre* mice (**Figure 5E-F**). In contrast, depletion of *Prenyl-Cdc42* mRNA had no apparent effect on regeneration in *Khsrp^fl/fl^* mice, which have wild type axonal KHSRP levels (**Figure 5C; Suppl. Figure S4**). Taken together, these findings indicate that stabilization of *Prenyl-Cdc42* mRNA in sciatic nerve axons contributes to the accelerated regeneration that we previously reported in KHSRP knockout mice.

### CSPG exposure decreases axonal Prenyl-Cdc42 mRNA by increasing axonal KHSRP

Since CSPG exposure also decreased axonal *Prenyl-Cdc42* mRNA (see Figures 1-2), we asked if the decrease in *Prenyl-Cdc42* mRNA after aggrecan exposure is mediated by KHSRP. Axons of *Khsrp^+/+^* neurons showed the anticipated decline in axonal *Prenyl-Cdc42* smFISH signals following aggrecan exposure; however, there was no change in the axonal *Prenyl-Cdc42* mRNA smFISH signals in *Khsrp^-/-^* neurons following aggrecan exposure (**Figure 6A-B; Suppl. Figure S5A**). Since axonal translation of *Khsrp* mRNA is activated by increased axoplasmic Ca^2+^ (Patel et al., 2022) and CSPGs are known to increase axonal Ca^2+^ (Kalinski et al., 2019; Snow et al., 1994), we asked if Ca^2+^ is necessary for the aggrecan-induced decrease in axonal *Prenyl-Cdc42* mRNA. Chelation of intracellular Ca^2+^ using BAPTA-AM blocked the aggrecan-induced decrease in axonal *Prenyl-Cdc42* mRNA (**Figure 6C-D; Suppl. Figure S5B**). Aggrecan treatment also selectively increased axonal and not cell body KHSRP levels (**Figure 6E-F; Suppl. Figure S5C**). Taken together, these studies indicate that KHSRP binds to the 3’UTR AU-rich region within nt 801-875 of *Prenyl-Cdc42* mRNA, which is functionally distinct from its axonal localization motif (nt 764-800), and Ca^2+^-dependent elevation of axonal KHSRP promotes decay of axonal *Prenyl-Cdc42* mRNA to slow axon growth.

**Figure 6:**
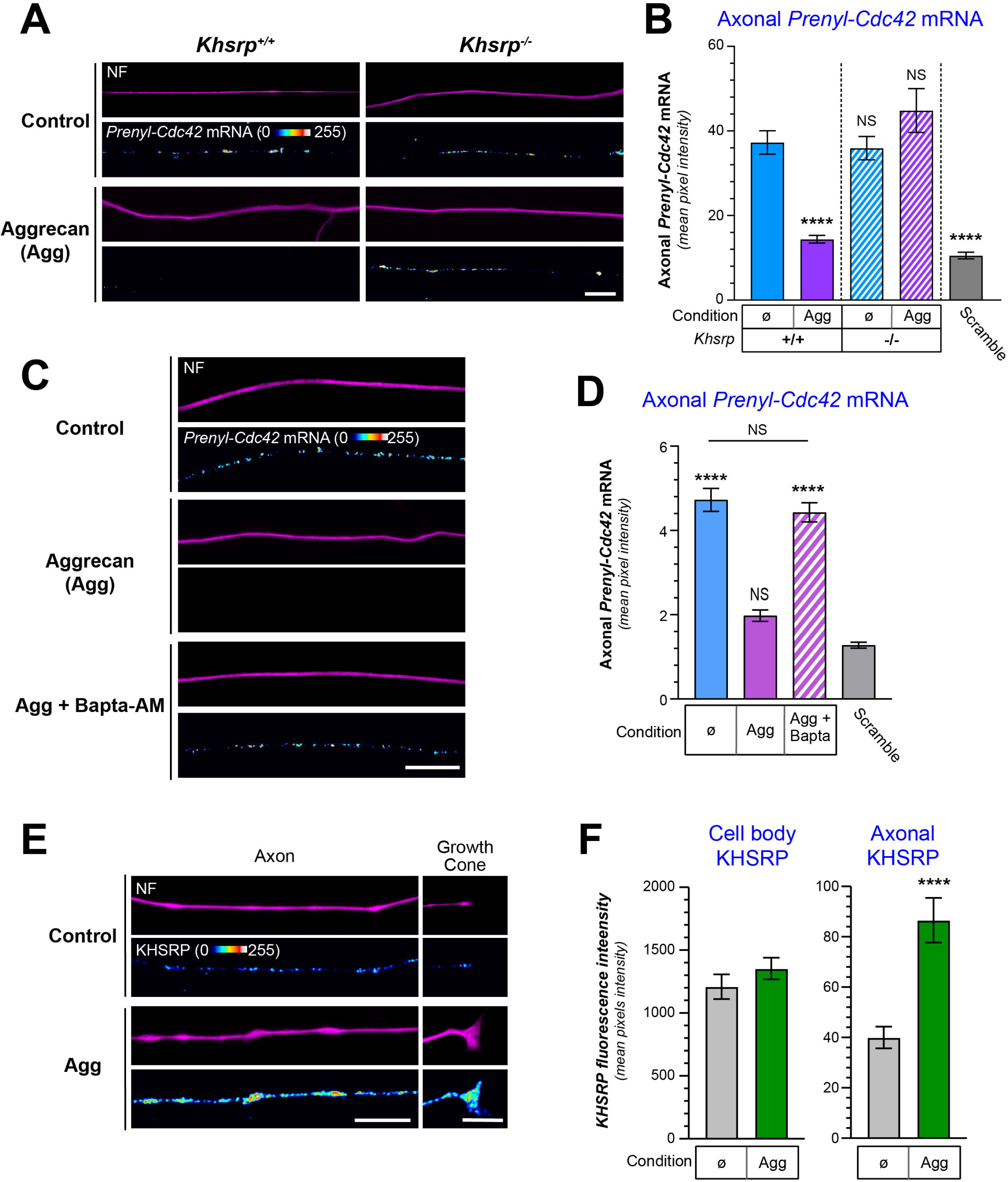
Aggrecan-mediated increase in axonal KHSRP depletes axonal Prenyl-Cdc42 mRNA. **A)** Representative exposure-matched smFISH and IF images for *Prenyl-Cdc42* mRNA and NF in adult mouse *Khsrp^+/+^* or *Khsrp^-/-^* DRG neuron cultures treated with 50 ng/ml aggrecan; see Suppl. Figure S5A for representative images of scrambled smFISH probe [Scale bar = 10 µm]. **B)** Quantitation of smFISH signal intensities shown as mean pixel intensity above background ± SEM for axons (N ≥ 40 neurons in three independent cultures; **** P <0.001 as compared to untreated KHSRP^+/+^ by one-way ANOVA with pair-wise comparison with Tukey post-hoc tests). **C)** Representative exposure-matched smFISH and IF images for *Prenyl-Cdc42* mRNA plus NF in adult DRG neuron cultures treated with 50 ng/ml aggrecan ± 3 µM BAPTA-AM; see Suppl. Figure S5B for representative images of scrambled smFISH probe [Scale bar = 10 µm]. **D)** Quantification of smFISH signal intensities shown as mean pixel intensity above background for axons ± SEM (N ≥ 40 neurons in three independent cultures; **** P < 0.001 compared to scramble probe as negative control by one-way ANOVA with pair-wise comparison with Tukey post-hoc tests) **E-F**) Representative exposure-matched IF images for KHSRP protein plus NF in adult DRG neuron cultures ± 50 ng/ml aggrecans are shown in E. Quantitation of cell body and axonal KHSRP signals from exposure-matched images is shown in F (see Suppl. Figure S6C for representative IF images for cell bodies; N ≥ 75 neurons over 3 separate cultures; **** P < 0.0001 by student’s t-test) [Scale bars = 10 µm].

## DISCUSSION

Axonally synthesized proteins have been shown to promote axon growth, axon survival, presynaptic plasticity, and injury signaling (Dalla Costa et al., 2020). Hundreds to thousands of mRNA are now known to localize into neuronal axons (Kar et al., 2018). *Trans-*acting proteins binding to *cis*- elements or motifs within the mRNAs, typically within the mRNA’s UTRs, are responsible for their subcellular localization, but also can impact the translation, storage and stability of those mRNAs once they arrive at their subcellular locale (Dalla Costa et al., 2020). There have not been any consensus sequence(s) identified that are shared across many axonal mRNAs bound by individual RNA binding proteins other than the ARE. Though not systematically analyzed in rodents, about 20% of human mRNAs are predicted to have 3’UTR AREs based on sequence analyses (Bakheet et al., 2018). We had previously reported that the 3’UTR of *Prenyl-Cdc42* mRNA is necessary and sufficient for its axonal localization (Lee et al., 2021) and we show here that the proximal region in *Prenyl-Cdc42* mRNA’s 3’UTR, nt 801-875, contains an AU-rich sequence indicative of an ARE. This region of *Prenyl-Cdc42* mRNA is greater than 85% conserved at primary sequence level across vertebrate orthologs. Sequence conservation in UTRs can point to functional regions of that RNA (Lee et al., 2018; Vuppalanchi et al., 2010), and we find that nt 764-800 and 801-875 constitute two distinct functional regions in *Prenyl-Cdc42* mRNA’s 3’UTR. The AU-rich region is within nt 801-875 motif and KHSRP binds to this motif confirming that it functions as an ARE-like motif, while nt 764-800 promotes axonal localization of *Prenyl-Cdc42* mRNA through an, as yet, unknown *trans*-acting protein(s).

ARE-binding has been demonstrated for many different RBPs, including KHSRP, HuC (ELAVL2), HuD (ELAVL4), HuR (ELAVL3), and hnRNPD (Bakheet et al., 2018). Of these, KHSRP and HuD localize to axons and have been suggested to compete for binding to overlapping ARE-containing mRNA populations (Gardiner et al., 2015). For example, the ARE in the 3’UTR of *Gap43* mRNA drives its localization into axons through HuD binding in a complex with Zip Code Binding Protein 1 (ZBP1) (Yoo et al., 2013). HuD binding also stabilizes *Gap43* mRNA (Gomes et al., 2017), with KHSRP binding through its KH4 domain promoting the *Gap43* mRNA’s decay (Bird et al., 2013)*. Neuritin* (*Nrn1*) mRNA also has a 3’UTR ARE motif that HuD binds in complex with SMN1 protein; this interaction is needed for its localization into axons of cortical but not sensory axons (Akten et al., 2011; Merianda et al., 2013). *Nrn1* mRNA localization into sensory axons requires a 5’UTR motif that hnRNP-H1, H2, and F bind (Lee et al., 2018; Merianda et al., 2013). *Prenyl-Cdc42* mRNA is similar to *Nrn1* mRNA’s behavior in sensory neurons, in that *Prenyl-Cdc42* mRNA’s localization motif is distinct from its ARE. It is not clear if HuD binds to *Prenyl-Cdc42*’s ARE, though HuD was shown to bind to *Cdc42* mRNA by RIP analyses using cDNA microarrays, that study did not distinguish *Prenyl-* and *Palm-Cdc42* mRNA isoforms (Bolognani et al., 2010). Nonetheless, it is clear that KHSRP binds to *Prenyl-Cdc42*’s AU rich motif from our data. Also, with the remarkable elevation of axonal *Prenyl-Cdc42* mRNA in the *Khsrp^-/-^* mice, KHSRP’s interaction with *Prenyl-Cdc42* mRNA depletes the transcript from distal axons. We previously showed that *Prenyl-Cdc42* mRNA level is very low in uninjured/non-growing axons (Lee et al., 2021). The *Khsrp^-/-^* mice showed increased in axonal *Prenyl-Cdc42* mRNA in uninjured conditions indicating that the mRNA is indeed transported into uninjured axons where basal levels of KHSRP likely promote its decay. Thus, KHSRP is likely utilized to dampen axonal synthesis of Prenyl-CDC42 to prevent or attenuate axon growth in uninjured axons and slow growth of regenerating axons. Consistent with this, depleting *Prenyl-Cdc42* mRNA from PNS axons prevents the accelerated nerve regeneration seen in *Khsrp^-/-^* mice, emphasizing the functional significance of KHSRP’s effect on axonal *Prenyl-Cdc42* mRNA.

KHSRP is a multifunctional RNA binding protein that has been implicated in RNA splicing, RNA transport and decay as well as microRNA biogenesis in addition to promoting ARE-containing mRNA decay (Briata et al., 2016). In previous studies, we had not been able to show any role for axonal KHSRP in microRNA biogenesis (Olguin et al., 2022). KSHRP has 4 KH RNA binding domains, with KH 1-2 domains needed for its RNA splicing function and KH 3-4 needed for its role in RNA decay promotion (Gherzi et al., 2004). Consistent with this, we previously showed that axonal levels of ARE-containing mRNAs are increased when KHSRP’s KH4 is deleted (Bird et al., 2013; Patel et al., 2022). Transcriptome analyses of *Khsrp^-/-^* mouse brain RNA combined with RIP-sequencing for KHSRP’s RNA interactome in wild type mouse brains showed that KHSRP binds to over 400 mRNA targets and promotes their decay (Olguin et al., 2022). *Cdc42 effector protein 3* mRNA was shown to increase in *Khsrp^-/-^* mouse brain, but *Cdc42* mRNA levels were not affected in those analyses of whole brain. However, both *Prenyl-Cdc42* and *Palm-Cdc42* mRNA isoforms were identified in KHSRP immunoprecipitates, showing that both isoforms are targets for regulation by KHSRP (Olguin et al., 2022). The discrepancy between KHSRP binding in wild type mice and lack of *Prenyl-Cdc42* mRNA elevation in cortical brain lysates of *Khsrp^-/-^* mice seen by Olguin et al. (2022) likely results from the selective interaction of KHSRP with axonal *Prenyl-Cdc42* mRNA that we report herein. This emphasizes that subcellular RNA-protein interactions and functional effects of those can be missed when looking at whole cell or tissue preparations.

The outcome of actin filament polymerization by CDC42 activation can be countered by actin filament depolymerization upon RHOA activation (Hall, 1998). Both CDC42 and RHOA must be activated by GTP binding (Hall, 1998). Differential regulation of axonal *Prenyl-Cdc42* and *RhoA* mRNA levels and translation in response to the growth-inhibiting CSPG but not the growth-promoting neurotrophin exposure suggest that post-transcriptional regulation of these Rho GTPases can impact axon growth. RHOA activation leads to growth cone collapse and axon retraction and inhibition of the RHOA/ROCK pathway supports axon growth on non-permissive substrates in cultured neurons, including the CSPG used here (Borisoff et al., 2003; Monnier et al., 2003). Traumatic CNS injury such as spinal cord injury (SCI) causes increased levels of growth-inhibiting molecules in the extracellular environment adjacent to the injury, which include CSPGs and myelin proteins (Quraishe et al., 2018). While extent of the contributions of these growth-inhibitory molecules to regeneration failure in the CNS brings some controversy (Sofroniew, 2018), blocking their effects has been proposed as neural repair strategy. RHOA inhibition has been tested pre-clinically and clinically as a therapeutic strategy to overcome the inhibitory environment of the injured CNS. A meta-analysis of experimental SCI models published over 2003-2018 showed that some but not all interventions to inhibit the RHOA pathway promoted *in vivo* axon regeneration (Luo et al., 2021). However, local delivery of the RHOA Inhibitor VX-210 did not prove effective for recovery in acute human cervical SCI (Fehlings et al., 2021), so it is unclear if other strategies to inhibit the RHOA pathway could bring effective *in vivo* SCI treatment options. Considering that CSPGs not only increase axonal RHOA but also deplete *Prenyl-Cdc42* mRNA from axons, our data raise the possibility that inhibition/inactivation of the RHOA pathway still leaves the axon in a low growth state since this would not prevent the depletion of *Prenyl-Cdc42* mRNA from axons in the injured CNS. Indeed strategies that increase axonal CDC42 activity may be needed to effectively promote axon regeneration in the non-permissive environment of the injured CNS.

CSPGs binding to the transmembrane receptors PTPσ and LAR activates RHOA/ROCK signaling to inhibit axon growth (Ohtake et al., 2016). *RhoA* mRNA was previously been shown to localize into axons (Wu et al., 2005), and its local translation was subsequently shown to be increased by CSPGs (Walker et al., 2012b). CSPG treatment increases intra-axonal Ca^2+^ in cultured DRG neurons (Snow et al., 1994). Translation of some mRNAs, including axonal *Khsrp* mRNA (Patel et al., 2022), is increased by Ca^2+^- dependent activation of PERK and subsequent phosphorylation of eIF2α (Boye and Grallert, 2020). Thus, a CSPG-driven increase in axonal Ca^2+^ could indeed increase local *Khsrp* mRNA translation to subsequently promote intra-axonal decay of *Prenyl-Cdc42* mRNA. Consistent with this, the CSPG- dependent depletion of *Prenyl-Cdc42* mRNA from axons was attenuated by chelating intra-cellular Ca^2+^ with BAPTA-AM. CSPGs as well as the CNS axon growth-inhibiting myelin-associated glycoprotein (MAG) attenuate axonal transport of mitochondria through a mechanism requiring elevation of axonal Ca^2+^ and activation of RHOA (Kalinski et al., 2019). This raises the possibility that signals from other CNS growth-inhibitory molecules similarly bring a dual hit to block axon regeneration by decreasing Prenyl-CDC42 and increasing RHOA syntheses in distal axons. Given the reciprocal regulation of *RhoA* and *Prenyl-Cdc42* mRNAs by CSPGs and the increased axonal transport and translation of *Prenyl-Cdc42* mRNA in response to neurotrophins, optimal growth of injured axons in the CNS may require simultaneously inhibiting the RHOA pathway and activating axonal translation of *Prenyl-Cdc42* mRNA.

## RESOURCE AVAILABILITY

The authors will distribute materials, reagents, and protocols to qualified researchers in a timely manner. All unique/stable reagents generated in this study are available from the lead contact with a completed materials transfer agreement.

## ACKNOWLEDGEMENTS

This work was supported by grants from the NIH (R01-NS089633 to JLT and NPB; R01-GM146257 to QL and JLT; R21-NS133477 to JML) and the Dr. Miriam and Sheldon G. Adelson Medical Research Foundation. JLT is the incipient University of South Carolina SmartState Chair in Childhood Neurotherapeutics, QL is the incipient University of South Carolina SmartState Chair in Neurotherapeutics, and JML is the incipient Carol and John Cromer ’63 Family Endowed Professor at Clemson University.

## AUTHOR CONTRIBUTIONS

MDZ, LSV, SJL, JML, and JLT designed experiments.

MDZ, LSV, SM, KT, AL, and SJL performed experiments.

MDZ, LSV, SM, KT, NPB, JML, and JLT provided data analyses.

LSV, JML, and JLT supervised the work.

MDZ, NPB, QL, JML, and JLT wrote and edited the manuscript.

NPB, QL, JML, and JLT obtained grants funding the work.

## DECLARATION OF AI AND AI-ASSISTIVE TECHNOLOGIES

Not applicable.

## SUPPLEMENTAL INFORMATION

STAR Methods

Supplemental Figures + Supplemental Figure Legends (5)

## SUPPLEMENTAL FIGURE LEGENDS

**<u>Supplemental Figure S1:</u> *Differential regulation of axonal Prenyl-Cdc42 and RhoA mRNA levels and translation (accompanying Figure 1-2).***

**A)** Representative IF images with no primary antibody as negative control for Figure 1D (see Figure 1E-F for quantifications).

**B-C)** Representative FRAP image sequences for DRG neurons co-transfected with GFP^MYR^5’/3’Cdc42 (B), and mCherry^MYR^5’/3’RhoA (C) at 72 h post-transfection are shown. Boxed regions represent the photobleached ROIs (see quantification in Figure 2B-C) [Scale bar = 20 µm].

**<u>Supplemental Figure S2:</u> *Sequence alignment for vertebrate Prenyl-Cdc42 mRNA orthologs.***

*Clustal Omega* multiple sequence alignment (Sievers et al., 2011) for the 3’UTR of *Prenyl-Cdc42* mRNAs are shown. Blue boxed regions show nucleotide conservation across orthologs. Nucleotide numbers labelled above start at the first nucleotide of the 3’UTR for *Xenopus tropicalis*. Beneath are graphical representations of consensus (% identity) and occupancy as well as a consensus aligned sequence.

**<u>Supplemental Figure S3:</u> *Cell body expression of GFP^MYR^3’prenyl-cdc42 mRNAs (accompanies data in Figure 3).***

**A)** Representative smFISH images for scramble FISH probe as negative control exposure matched to those in Figure 3B (see Figure 3D for quantitative data) [Scale bar = 10 µm].

**B-C**) Quantitation of smFISH signal intensities shown as mean pixel intensity above background ± SEM for cell bodies corresponding to Figures 3D-E (*N* ≥ 40 neurons in three independent cultures; NS = not significant between indicated data pairs by one-way ANOVA, pair-wise comparison with Tukey post-hoc tests).

**<u>Supplemental Figure S4:</u> *KHSRP regulates axonal Prenyl-Cdc42 mRNA levels (accompanies data in Figures 4-5).***

**A)** Quantitation of smFISH signal intensities shown as mean pixel intensity above background ± SEM for cell bodies (see Figure 4D for representative images; *N* ≥ 30 neurons in three independent cultures; NS = not significant between indicated data pairs by one-way ANOVA with pair-wise comparison with Tukey post-hoc tests).

**B)** Representative smFISH images for 7 day post-crush injured sciatic nerve showing signals for scramble probe exposure matched to Figure 5A (see Figure 5B for quantitation) [Scale bar = 5 µm].

**C)** RTddPCR for sciatic nerve axoplasm for *Prenyl-Cdc42* mRNA in sciatic nerve axoplasm from *Khsrp^-/-^* mice treated with siRNA polymersomes (NP-siRNA) targeting *Prenyl-Cdc42* mRNA vs. scrambled siRNA sequences at 7 d post-crush (Con; N = 6 mice; NS = not significant, **** p < 0.001 by students T-test).

**D-E)** Representative IF images for SCG10 in sciatic nerves of *Khsrp^fl/fl^* mice (D) at 14 d post-crush with NP-siRNA injection at 7 d post crush as in Figure 5 E-F. Quantitation of regeneration indices shown as in E (N = 3 animals per condition; * P < 0.05 by repeated measures ANOVA) [Scale bar = 100 µm].

**<u>Supplemental Figure S5:</u> *Aggrecan triggered increase in KHSRP is limited to axons (accompanies data from Figure 6)***

**A)** Representative smFISH images using scramble probe as negative control in adult mouse *Khsrp^-/-^* DRG neuron cultures exposure matched to Figure 6A (see Figure 6B for quantitation) [Scale bar = 10 µm]

**B)** Representative smFISH images using scramble probe as negative control in adult mouse DRG neuron cultures exposure matched to Figure 6C (see Figure 6D for quantitation) [Scale bar = 10 µm].

**C)** Representative exposure-matched IF images for KHSRP protein in cell bodies of cultured mouse DRG neurons exposed to aggrecan as in Figure 6E (see Figure 6F for quantitation) [Scale bar = 25 µm].

## STAR METHODS

### KEY RESOURCES TABLE

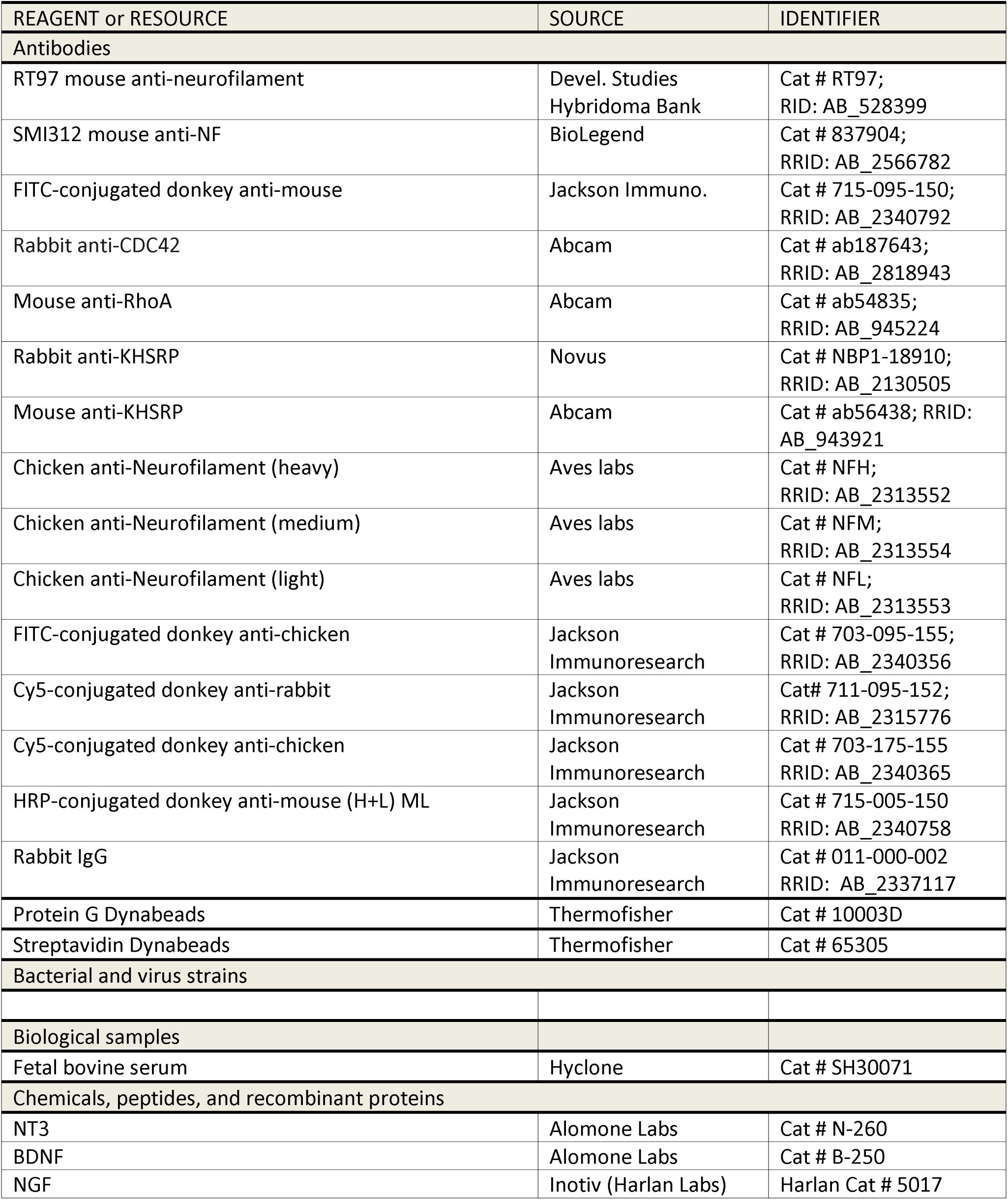
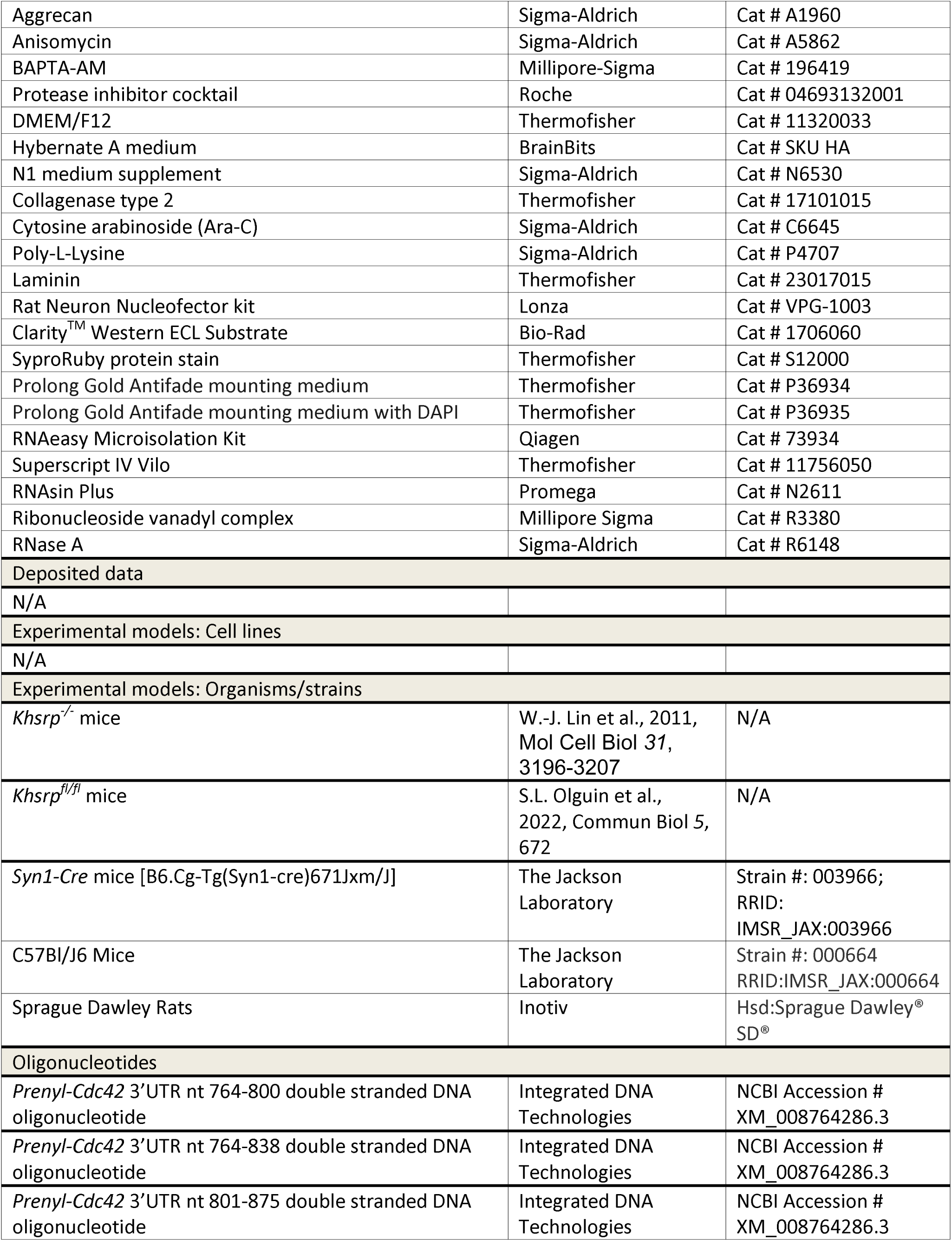
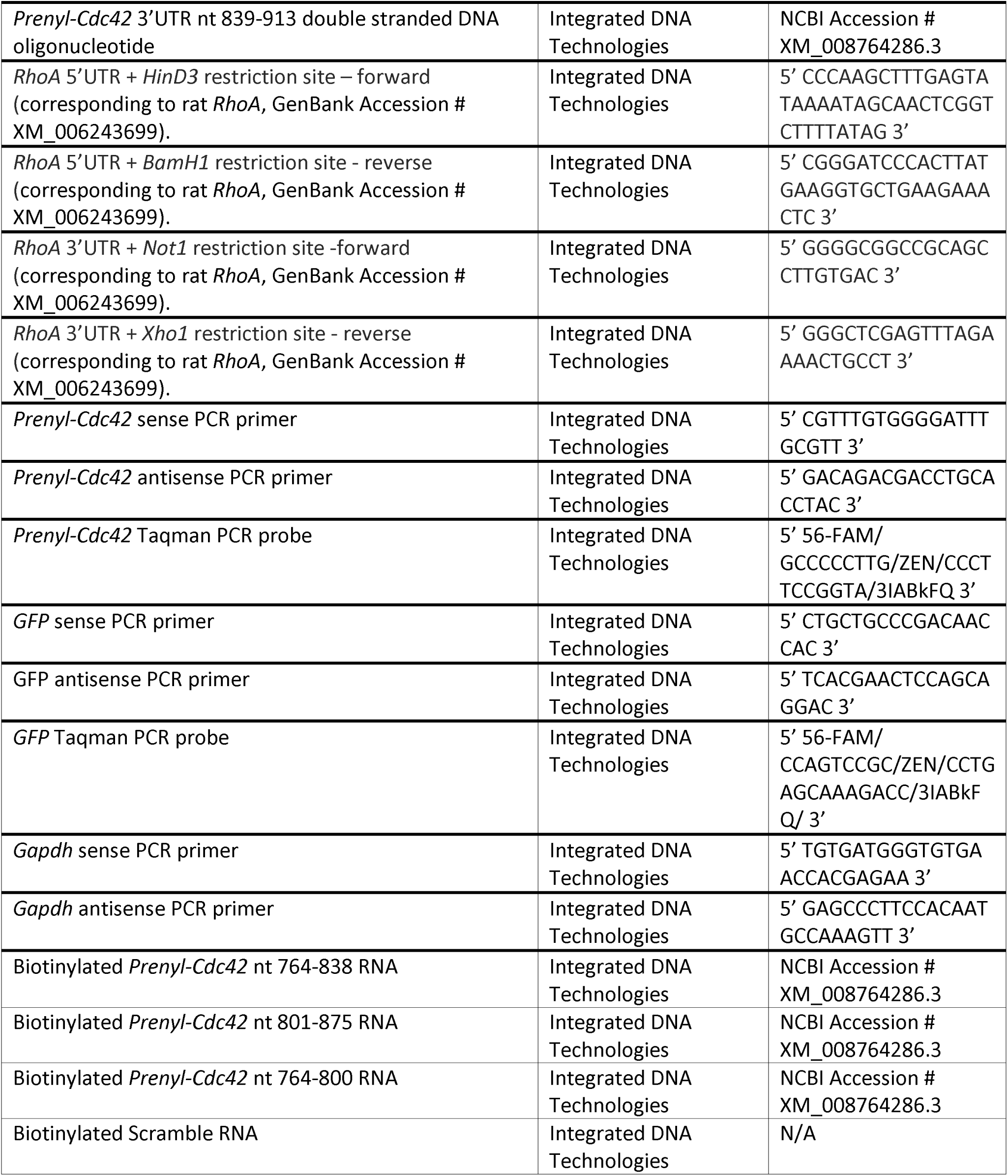
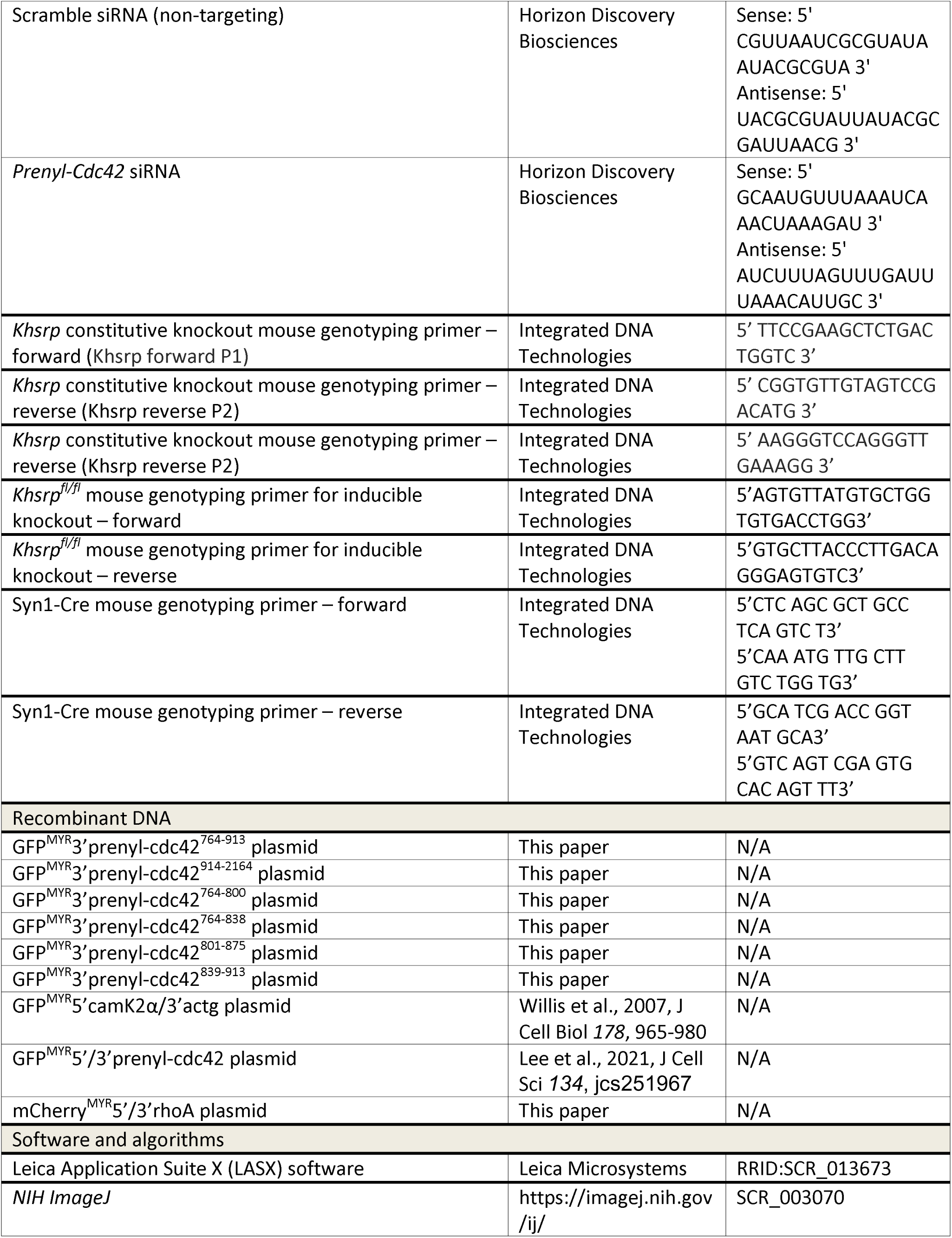
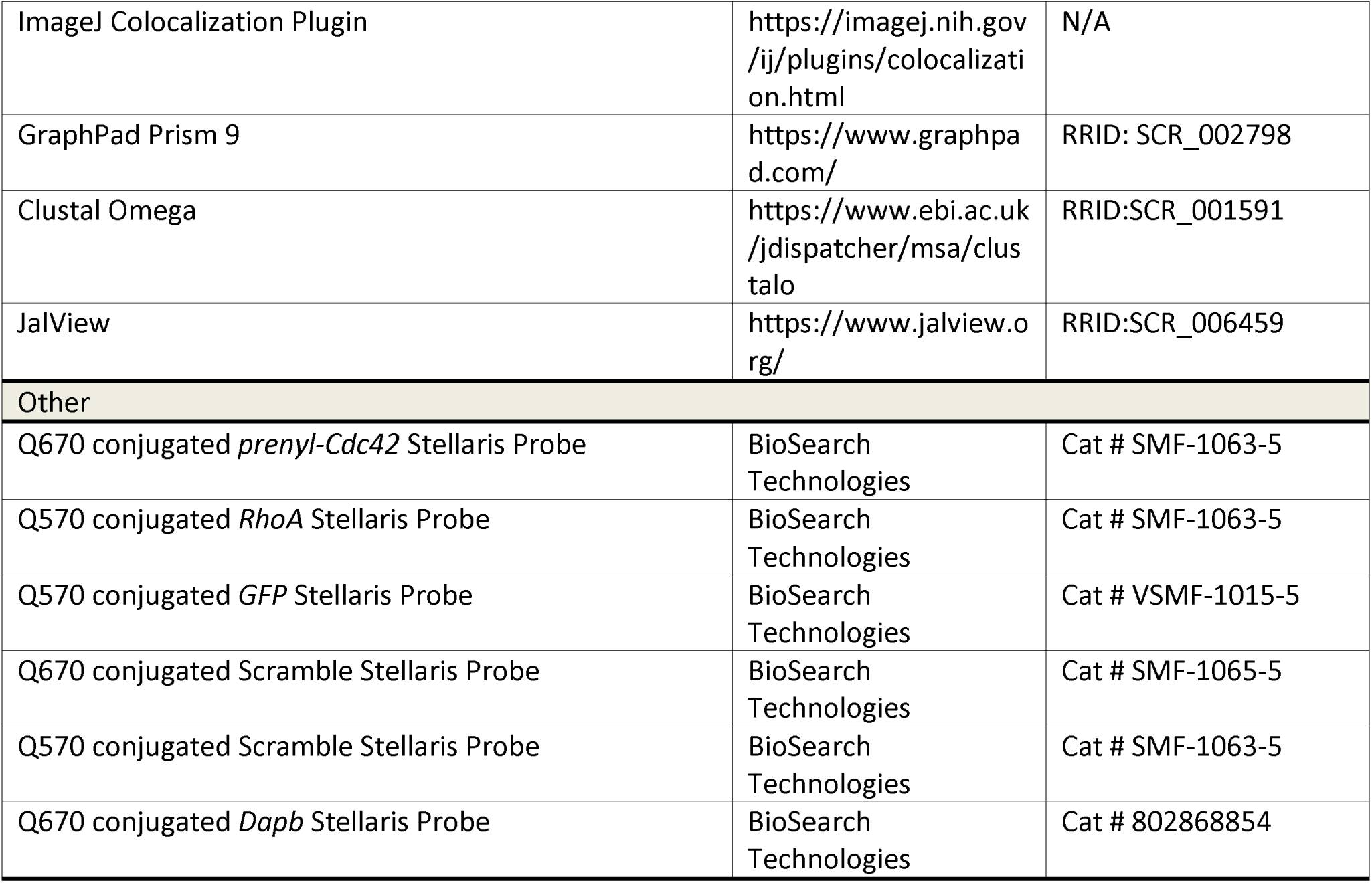

### CONTACT FOR REAGENT AND RESOURCE SHARING

Jeff Twiss

Dept Biological Sciences

715 Sumter Street, CLS 401

University of South Carolina

Columbia, SC 29204 USA

phone – 803-777-9215

email –

### EXPERIMENTAL MODEL AND SUBJECT DETAILS

#### Animal care and use

Institutional Animal Care and Use Committee of the University of South Carolina approved all animal procedures. Sprague Dawley rats (175-250 g) were used for preparing sciatic nerve axoplasm. Male and female wild type C57Bl/6 (*Khsrp^+/+^*), constitutive *Khsrp* knockout (*Khsrp^-/-^*) (Lin et al., 2011), and conditional *Khsrp^fl/fl^* (Olguin et al., 2022) mice were used for sciatic nerve injury and DRG culture experiments as indicated in the results. For neuronal specific *Khsrp* knockout, male *Khsrp^fl/fl^* mice were crossed to female B6.Cg-Tg(Syn1-cre)671Jxm/J (*Syn1-Cre*; Jackson Laboratories) mice. All animals were euthanized by CO_2_ asphyxiation per IACUC guidelines.

For nerve crush surgery, animals were anesthetized with isoflurane by inhalation (5% induction and 2% maintenance). Anesthetized animals were subjected to sciatic nerve crush at mid-thigh level as previously described (Twiss et al., 2000). Briefly, the nerve was exposed by blunt dissection and then crushed with # 2 fine jeweler’s forceps, twice for 15 sec each; success of the axotomy was monitored by the initial contraction of the hind limb upon applying pressure to the nerve and then lack of hind paw extension during and upon recovery from anesthesia.

For *in vivo* RNA depletion from sciatic nerve axons, polymersomes with siRNAs (see below) were delivered by injecting 6 µl of polymersome solution in 1x PBS that contained an equivalent of 40 nM siRNAs. Polymersomes were injected into the sciatic nerve of anesthetized mice (see above) at 7 days following nerve crush injury (performed as above) just proximal to the injury site. Delivery of polymersomes was confirmed by RTddPCR for *Prenyl-Cdc42* and *Gapdh* mRNAs, visualization of the DiD fluorophore in the nerve, and smFISH/IF for *Prenyl-Cdc42* mRNA and neurofilament protein.

### METHOD DETAILS

Mouse genotyping – Genotyping for constitutive Khsrp knockout was performed using PCR with primers spanning the exon 1 to exon 13 deletion of the mouse Khsrp gene or wild type sequence as previously described (Olguin et al., 2022). For this, DNA was extracted from ear punches taken at weaning. Primers used for genotyping are as follows (5’ to 3’): Khsrp forward P1 – TTCCGAAGCTCTGACTGGTC, Khsrp reverse P2 – CGGTGTTGTAGTCCGACATG, and Khsrp reverse P3 – AAGGGTCCAGGGTTGAAAGG. PCR products were analyzed by agarose gel electrophoresis with SYBR™ Safe DNA Gel Stain (Thermofisher). The Cre-loxP system is used to generate a conditional knockout of *Khsrp*. *Khsrp^fl/fl^* mice were generated by Biocytogen using CRISPR/EGE™-based gene editing to insert loxP sites between exons 1 and 2 and exons 6 and 7 that would result in a frameshift upon Cre-driven recombination but were not predicted to affect splicing of the *Khsrp* RNA transcript prior to any recombination (Patel et al., 2022). Genotyping for loxP insertion was performed using following primers (5’ to 3’): 5’ LoxP forward – AGTGTTATGTGCTGGTGTGACCTGG, 5’ LoxP reverse – GTGCTTACCCTTGACAGGGAGTGTC, 3’ LoxP forward – CTATGGTGTCACCTCTCAGTGCTGC, and 3’ LoxP reverse – CACGTAGAGGCCAAAGCAAGAGGAC. PCR products were analyzed by agarose gel electrophoresis with SYBR™ Safe DNA Gel Stain (Thermofisher). For specific Cre expression in neuronal cells, Syn1-Cre mice were used, with Cre expression driven by a rat Synapsin I promoter. The following primers were used to detect Syn1-Cre-mediated recombination (5’ to 3’): forward transgene – CTCAGCGCTGCCTCAGTCT, reverse transgene – GCATCGACCGGTAATGCA, forward IPC – CAAATGTTGCTTGTCTGGTG, and reverse IPC – GTCAGTCGAGTGCACAGTTT. PCR products were analyzed by agarose gel electrophoresis with SYBR™ Safe DNA Gel Stain (Thermofisher).

#### Primary neuron culture

Dissociated cultures of adult DRGs were prepared as described (Twiss et al., 2000). DRGs were harvested in Hybernate-A medium (BrainBits) and then dissociated with 2,000 units/ml Collagenase type 2 (Thermofisher) at 37°C, 5% CO_2_ for 15 min. Ganglia were triturated using a fire polished Pasteur pipet to fully dissociate, and then diluted in 9 volumes DMEM/F12 (Thermofisher) pelleted at 100 xg for 5 min. After pelleting, dissociated ganglia were washed in DMEM/F12 and then cultured in DMEM/F12, 1 x N1 supplement (Sigma-Aldrich), 10% fetal bovine serum (Hyclone), and 10 µM cytosine arabinoside (Sigma-Aldrich). For imaging experiments, dissociated DRGs were cultured on poly-L-lysine (Sigma-Aldrich) and laminin (Thermofisher)-coated glass coverslips.

For transfections, dissociated ganglia were pelleted at 100 x g for 5 min and resuspended in 100 µl ‘Nucleofector solution’ (Rat Neuron Nucleofector kit; Lonza). 4-6 µg of each plasmid was electroporated using the AMAXA Nucleofector device (G013 program; Lonza) before plating. Dissociated ganglia were then plated as above and analyzed 48-72 h later.

#### Plasmid constructs

Mammalian expression plasmids with the coding sequence of the fluorescent reporter enhanced green fluorescent protein (eGFP) containing cDNA corresponding to the 5’ and 3’UTRs of rat *Prenyl-Cdc42* (Lee et al., 2021) were used as a basis for 3’UTR deletion constructs GFP^MYR^3’prenyl-Cdc42^764-913^ and GFP^MYR^3’prenyl-Cdc42^914-2164^. Constructs were produced by digestion with either Not1 and BstX1 for GFP^MYR^3’prenyl-Cdc42^764-91*3*^, or Bstx1 and EcoR1 for GFP^MYR^3’prenyl-Cdc42^914-2164^ corresponding to rat *Prenyl-Cdc42* nt 764-913 or 914-2164 (GenBank Accession # XM_008764286). 3’ overhangs were then filled using Klenow fragment (New England Biolabs) and re-ligated. The GFP coding sequence in these included a myristylation sequence (GFP^MYR^; originally provided by Dr. Erin Schuman, Max-Plank Inst., Frankfurt) (Aakalu et al., 2001) that limits diffusion of the GFP translation product.

To create expression constructs containing shortened portions of Cdc42 3’UTR, double stranded DNA segments, corresponding to nt 764-800, 764-838, 801-875, and 839-913 of rat *Prenyl-Cdc42* mRNA, were custom synthesized by Integrated DNA Technologies. These 3’UTR segments were engineered with 5’ Not1 and 3’ Xho1 restriction sites and used to replace the 3’UTR in GFP^MYR^5’CamK2α/3’actg plasmid. This plasmid contains the 5’UTR of calcium/calmodulin dependent protein kinase II alpha (CamK2α) that has previously been shown to lack any activity for axonal localization (Willis et al., 2007).

GFP^MYR^5’/3’prenyl-Cdc42 used in our FRAP analyses was previously generated in our lab (Lee et al., 2021). mCherry^MYR^5’/3’rhoA was generated by replacing the 5’ and 3’UTR of mCherry^MYR^511/311kpnb1 (Sahoo et al., 2020) with PCR generated sequences. Primers used for cloning are as follows (5’ to 3’): *RhoA* 5’UTR *HinD3* forward – CCCAAGCTTTGAGTATAAAATAGCAACTCGGTCTTTTATAG, *RhoA* 5’UTR *BamH1* reverse – CGGGATCCCACTTATGAAGGTGCTGAAGAAACTC, *RhoA* 3’UTR *Not1* forward – GGGGCGGCCGCAGCCTTGTGAC, *RhoA* 3’UTR *Xho1* reverse – GGGCTCGAGTTTAGAAAACTGCCT, corresponding to rat *RhoA* (GenBank Accession # XM_006243699). The 5’UTR was engineered with 5’ Hind3 and 3’ BamH1 restriction sites and the 3’ UTR was engineered with 5’ Not1 and 3’ Xho1 restriction sites and used to replace the 5’ and 3’ UTRs of mCherry^MYR^5’/3’kpnb1 (Sahoo et al., 2020).

#### Polymersome delivery of siRNAs

Synthetic siRNAs were purchased from Dharmacon (Horizon Discovery Biosciences). The *Prenyl-Cdc42* and non-targeting control siRNA sequences were previously published (Lee et al., 2021). siRNAs were first tested for specificity in primary mouse dissociated DRG cultures using Dharmafect 3 (Horizon Discovery Biosciences) for transfections and RTddPCR to validate depletion of *Prenyl-Cdc42* mRNA in mouse DRG cultures. Validated siRNAs were then packaged into polymeric nanoparticles called polymersomes that were labeled with a peptide of rabies virus glycoprotein, RVG29 (called RVG herein). Polymersomes were made from solvent injection of a 50:50 mixture of block co-polymer polyethylene glycol (PEG, 1000 kDa)-b-polylactic acid (PLA, 5000 kDa) and PEG(1000)-b-PLA(5000)-maleimide, then lyophilized prior to siRNA encapsulation. Cysteine conjugated RVG is added to solution, enabling a thiol coupling reaction on the polymersome surface to attach the RVG. RVG attachment is confirmed via a shift in surface charge from negative to positive via zeta potential measurements (Trumbull et al., 2024). RVG-tagged polymersomes co-encapsulated 13.6 ± 4.6 µg siRNA/mg polymer with 53 ± 8 µg membranous DiD/mg polymer. RVG-tagged siRNA loaded polymersomes were concentrated to 100 mM prior to injection.

#### Fluorescence in situ hybridization and immunofluorescence

Custom Stellaris oligonucleotide probes with 511 Cy3 or Cy5 labels (BioSearch Tech.) were used for smFISH combined with IF to detect endogenous *Cdc42, RhoA* and *GFP* mRNAs (probe sequences available upon request). Scrambled probes were used as control for specificity; samples processed without the addition of primary antibody were used as control for antibody specificity. For cultured neurons, coverslips with dissociated DRGs were briefly rinsed in phosphate buffered saline (PBS), fixed for 15 min in 2% PFA in PBS, and then processed for pre-hybridization and hybridization as described (Kalinski et al., 2015). Primary antibodies to detect neurofilament (NF) consisted of a cocktail of mouse RT97 (1:500, Devel. Studies Hybridoma Bank) and SMI312 (1:250; BioLegend). FITC-conjugated donkey anti-mouse (1:200, Jackson ImmunoRes.) was used as the secondary antibody.

smFISH/IF was performed on sciatic nerve cryosections as described previously (Kalinski et al., 2015). Briefly, sciatic nerve segments were immersion fixed overnight in 2% PFA in PBS at 4°C and then cryoprotected overnight in 30% sucrose in 1x PBS at 4°C. Samples were processed for cryosectioning at 10 µm thickness, sections were adhered to Superfrost^plus^ glass slides (Fisher) and then stored at −20°C until use. Slides were dried at 37°C for 1 h and then brought to room temperature. All subsequent steps performed at room temperature unless indicated otherwise. Sections were washed for 10 min in PBS, then 10 min in 20 mM glycine three times followed by three 5 min incubations in fresh 0.25 M NaBH_4_. Sections were rinsed in 0.1 M triethanolamine (TEA), incubated for 10 min in 0.25% acetic anhydride in 0.1 M TEA, and washed twice with 2x saline-sodium citrate (SSC) buffer. Sections were then dehydrated through graded ethanol solutions (70, 95, and 100 % for 3 min each) followed by delipidation in chloroform for 5 min. Sections were rehydrated in 100 and 95% ethanol for 3 min each, equilibrated in 2x SSC and then incubated at 37°C in a humidified chamber in hybridization buffer (50 % dextran sulphate, 10 μg/ml E. coli tRNA, 10 mM ribonucleoside vanadyl complex [Millipore Sigma], 80 μg BSA, and 10 % formamide in 2× SSC) for 5 min. Hybridization and immunolabeling were performed overnight in a humidified chamber in hybridization buffer containing 7 µM of each probe and RT97 antibody (1:100). Sections were then washed twice in 2x SSC plus 10 % formamide at 37°C for 30 min and once in 2x SSC for 5 min. After permeabilization in PBS plus 1 % Triton X-100 (PBST) for 5 min, sections were incubated for 1 h in FITC-conjugated donkey anti-mouse-IgG antibody (1:200) in 0.3 % Triton X-100 supplemented with 1× blocking buffer (Roche). After washing in PBS for 5 min, sections were post-fixed in 2% PFA in PBS for 15 min (Spillane et al., 2013), washed in PBS three times for 5 min, and then rinsed in DEPC-treated water. Both the cultured neurons and tissues were mounted using Prolong Gold Antifade (Invitrogen).

All smFISH/IF performed on tissue sections was imaged by confocal microscopy using a Leica SP8X confocal microscope with HyD detectors and matched post-processing measures to distinguish axonal signals from non-neuronal signals. Scrambled probes were used to assign maximum acquisition parameters to limit any nonspecific signal from the probes.

Standard IF was performed as previously described (Gomes et al., 2017) with all steps at room temperature unless specified otherwise. Coverslips were fixed with 4% PFA in PBS for 15 min and washed 3 times in PBS. PBS washed neurons were permeabilized with 0.3 % Triton X-100 in PBS for 15 min and then blocked in PBST plus 5 % BSA for 1 h. Neurons were incubated with primary antibodies overnight at 4°C. Primary antibodies consisted of rabbit anti-CDC42 (1:500; Abcam), mouse anti-RhoA (1:25; Abcam), and chicken anti-NF (1:500, NFM, NFL and NFH, Aves labs). After washing in PBS, coverslips were incubated with combination of FITC-conjugated donkey anti-chicken and Cy5-conjugated donkey anti-rabbit or anti-chicken (all at 1:500; Jackson ImmunoRes.) as secondary antibodies for 1 h. After 1 h, coverslips were washed 3 times in PBS, rinsed with distilled H_2_O, and mounted with Prolong Gold Antifade with DAPI (Invitrogen).

#### Fluorescence recovery after photobleaching

FRAP analyses were performed as published with minor modifications (Vuppalanchi et al., 2010). DRG neurons were transfected with GFP^MYR^5’/3’prenyl-Cdc42 and mCherry^MYR^5’/3’RhoA as above. Cells were maintained at 37°C, 5 % CO_2_ during imaging. 488 nm and 587 nm laser lines on a Leica SP8X confocal microscope with HyD detectors were used to bleach GFP and mCherry signal, respectively (argon laser at 70 % power, pulsed every 0.82 sec for 80 frames). Leica 63x/1.4 NA oil immersion objective was used with the confocal pinhole set to 3 Airy units to ensure full thickness bleaching and acquisition (Yudin et al., 2008). 488 nm and 587 nm laser lines on the Leica SP8X confocal microscope with HyD detectors were used to bleach GFP and mCherry signals, respectively (argon laser at 70 % power, pulsed every 0.82 sec for 80 frames). Prior to photobleaching, two frames were acquired at 60 sec intervals to determine baseline fluorescence in the region of interest (ROI; 15 % laser power, 498-530 nm for GFP; 597-630 nm for mCherry). The same excitation and emission parameters were used to assess recovery over 15 min post-bleach with images acquired every 30 sec. For some experiments, 50 µg/ml of aggrecan (Sigma-Aldrich) or a neurotrophin cocktail consisting of neurotrophin 3 (NT3; Alamone Labs), brain-derived neurotrophic factor (BDNF; Alamone Labs) and NGF (10 µg/ml each; Inotiv) was bath applied immediately prior to imaging. To test whether any fluorescence recovery in axons was due to translation, DRG cultures were treated with 100 µM anisomycin (Sigma) for 30 min prior to photobleaching.

#### RNA isolation and analyses

RNA was isolated from mouse DRG culture lysates and affinity pull down samples using the RNeasy Microisolation kit (Qiagen). Fluorimetry with Ribogreen (Invitrogen) was used for RNA quantification for total RNA isolates. For analyses of total RNA levels and inputs for co-immunoprecipitation, RNA yields were normalized for mass across samples prior to reverse transcription (RT) using Superscript IV Vilo (Thermofisher). For co-immunoprecipitation of RNA, samples were processed based on equivalent proportions of the precipitate rather than normalizing to RNA mass. Droplet digital PCR (ddPCR) was performed using Taqman probe sets (Integrated DNA Tech) and QX200^TM^ droplet reader (Bio-Rad). Primer probe sets were as follows (5’ to 3’): prenyl-Cdc42 sense primer – CGTTTGTGGGGATTTGCGTT, prenyl-Cdc42 antisense primer – GACAGACGACCTGCACCTAC, prenyl-Cdc42 Probe - /56-FAM/GCCCCCTTG/ZEN/CCCTTCCGGTA/3IABkFQ/, GFP sense primer – CTGCTGCCCGACAACCAC, GFP antisense primer – TCACGAACTCCAGCAGGAC, GFP probe - /56- FAM/CCAGTCCGC/ZEN/CCTGAGCAAAGACC/3IABkFQ/.

#### Affinity isolation of RNA-interacting proteins

RNA-Protein pull-down was performed as described (Doron-Mandel et al., 2016). We used axoplasm from sciatic nerve as a source of proteins. To obtain enriched axonal contents, approximately 3 cm segments of rat sciatic nerve were dissected and axoplasm was extruded into 20 mM HEPES [pH 7.3], 110 mM potassium acetate, and 5 mM magnesium acetate (nuclear transport buffer) supplemented with 1x protease/phosphatase inhibitor cocktail (Roche) and 40 U/µl RNasin Plus (Promega) as previously described (Hanz et al., 2003). After clearing by centrifugation at 20,000 xg at 4°C for 30 min, supernatants were mixed with 5’ biotin-conjugated RNA oligonucleotides (Integrated DNA Technologies), which had been adsorbed to streptavidin (SA) dynabeads (Thermofisher) (Lee et al., 2018), and incubated for 4 h at 4°C. Beads were precipitated using a magnetic rack and then washed extensively with 10 mM HEPES (pH 7.4), 3 mM MgCl_2_, 250 mM NaCl, 1 mM DTT, and 5% glycerol. Bound proteins were eluted by treating with 50 µg/ml RNase A (Sigma-Aldrich) in wash buffer for 15 min at 37°C (Lee et al., 2018). Proteins were denatured by boiling at 95°C in Laemmli buffer for 5 min, fractioned by SDS/PAGE, and then transferred to nitrocellulose membranes for Immunoblotting.

#### RNA co-immunoprecipitation

For co-precipitating RNAs with proteins, DRG cultures were lysed in 100 mM KCl, 5 mM MgCl_2_, 10 mM HEPES [pH 7.4], 1 mM DTT, and 0.5% NP-40 (‘RIP buffer’) supplemented with 1x protease inhibitor cocktail and RNasin Plus. Lysates were passed through 25 Ga needle 5-7 times and then cleared by centrifugation at 12,000 xg for 20 min. Cleared lysates were incubated with Protein G-Dynabeads (Thermofisher) for 30 min to reduce non-specific binding. After collection, supernatants were then incubated with rabbit anti-KHSRP (5 μg, Novus) or rabbit IgG (5 μg, Jackson ImmunoRes.) for 3 h at 4°C with rotation. Immunocomplexes were incubated with Protein G-Dynabeads for an additional 2 h at 4°C with rotation. Beads were washed six times with cold RIP buffer. An aliquot was reserved for validating protein precipitation (see below), and bound RNAs were isolated by addition of RNeasy Microisolation kit lysis buffer and analyzed by RTddPCR (see above).

For validation of pull-down of protein, the reserved aliquot from the immunoprecipitates was resuspended in 1 x Laemmli sample buffer and denatured by boiling at 95°C x 5 min. Supernatants were then processed for immunoblotting as described below, including a reserved aliquot from the input lysate for determining relative degree of RNA binding to KHSRP.

#### Protein electrophoresis and immunoblotting

Protein concentrations were determined by BCA assay. Cell lysates and axoplasm were normalized for concentration prior to electrophoresis or immunoprecipitation (see below). Lysates, immunoprecipitates, and RNA affinity isolates were denatured by boiling in Laemmli sample buffer, fractionated by SDS-PAGE, and electrophoretically transferred to nitrocellulose membranes. Blots were incubated for 1 h at room temperature in blocking buffer (5% non-fat dry milk in Tris-buffered saline with 0.1% Tween 20 [TBST]). Membranes were incubated overnight incubation at 4°C with rocking in mouse anti-KHSRP (1:1000; Abcam; Ab56438) diluted in TBST plus 5% BSA. After washing in TBST, blots were incubated HRP-conjugated anti-mouse IgG antibodies (1:5000; Jackson ImmunoRes.) diluted in blocking buffer for 1 h at room temperature. After washing in TBST (3 times), signals were detected using Clarity^TM^ Western ECL Substrate (Bio-Rad). ***In vitro oligo RNA-protein binding assay –*** 5’ biotin-conjugated RNA oligonucleotides were immobilized onto M-280 streptavidin (SA) dynabeads (Thermofisher) per manufacturer’s instructions. Equimolar amounts of purified recombinant KHSRP^FLAG^ protein and oligo-immobilized beads were incubated by rotation in 10 mM HEPES [pH 7.4], 3 mM MgCl_2_, 14 mM NaCl, 1 mM DTT, and 5% glycerol for 4 h at 4°C. Beads were precipitated using a magnetic rack and then washed extensively with 10 mM HEPES [pH 7.4], 3 mM MgCl_2_, 250 mM NaCl, 1 mM DTT, and 5% glycerol. Bound proteins were eluted by boiling at 95°C in Laemmli buffer for 5 min, fractioned by SDS/PAGE, and then transferred to PVDF membranes for Immunoblotting.

### QUANTITATION AND STATISTICAL DETAILS

All imaging experiments included at least three technical replicates for each culture, and each experiment was replicated across at least three separate culture preparations. For molecular studies using transfected cultures, analyses were performed across at least three separate culture preparations. Positive and negative controls were included as outlined above and in the Results section.

For smFISH on tissue sections the Z stacks of XY optical planes were captured at two locations along each nerve section. The *Colocalization Plug-in* for *NIH ImageJ* (https://imagej.nih.gov/ij/ plugins/colocalization.html) was used to extract RNA signals from smFISH probes in each optical plane that overlapped with NF signals as an ‘axon only’ mRNA signal. All smFISH signal quantifications for axonal mRNA signals from tissue sections were generated by analyses of pixel intensity across each XY plane of the extracted ‘axon only’ channels for the image sequences using ImageJ. These smFISH signal intensities across the individual XY planes were then normalized to the area of NF immunoreactivity in each XY plane and averaged across the image stack (Kalinski et al., 2015). The relative mRNA signal intensity was then determined from the average in each biological replicate.

For FRAP assays, fluorescent intensities in the bleached region of interest (ROI) were calculated using Leica LASX software. For normalizing across experiments, fluorescence intensity value for the ROI at t = 0 min postbleach from each image sequence was set as 0 %. The relative fluorescence recovery at each time point after photobleaching was then calculated by normalizing to the pre-bleach fluorescence intensity of the ROI (set at 100 %). All bleached ROIs were at least 250 µm from the cell soma, so any significant fluorescence recovery occurring in less than 15 min post-bleach interval can be attributed to local protein synthesis rather than anterograde transport of reporter protein into the ROI if recovery was significantly attenuated by protein synthesis inhibitors.

Quantitative data are reported as mean ± standard error of the mean (SEM) as indicated in the figure legends. Student’s *t*-test or one-way ANOVA with multiple comparisons and Tukey post-hoc analyses was used to test for significance between groups as indicated in figure legends. GraphPad Prism 9 software was used for all statistical analyses.

