## Supplementary figures and images for "KHSRP-mediated Decay of Axonally Localized Prenyl-Cdc42 mRNA Slows Nerve Regeneration"

### Supplemental Figures S1-S5

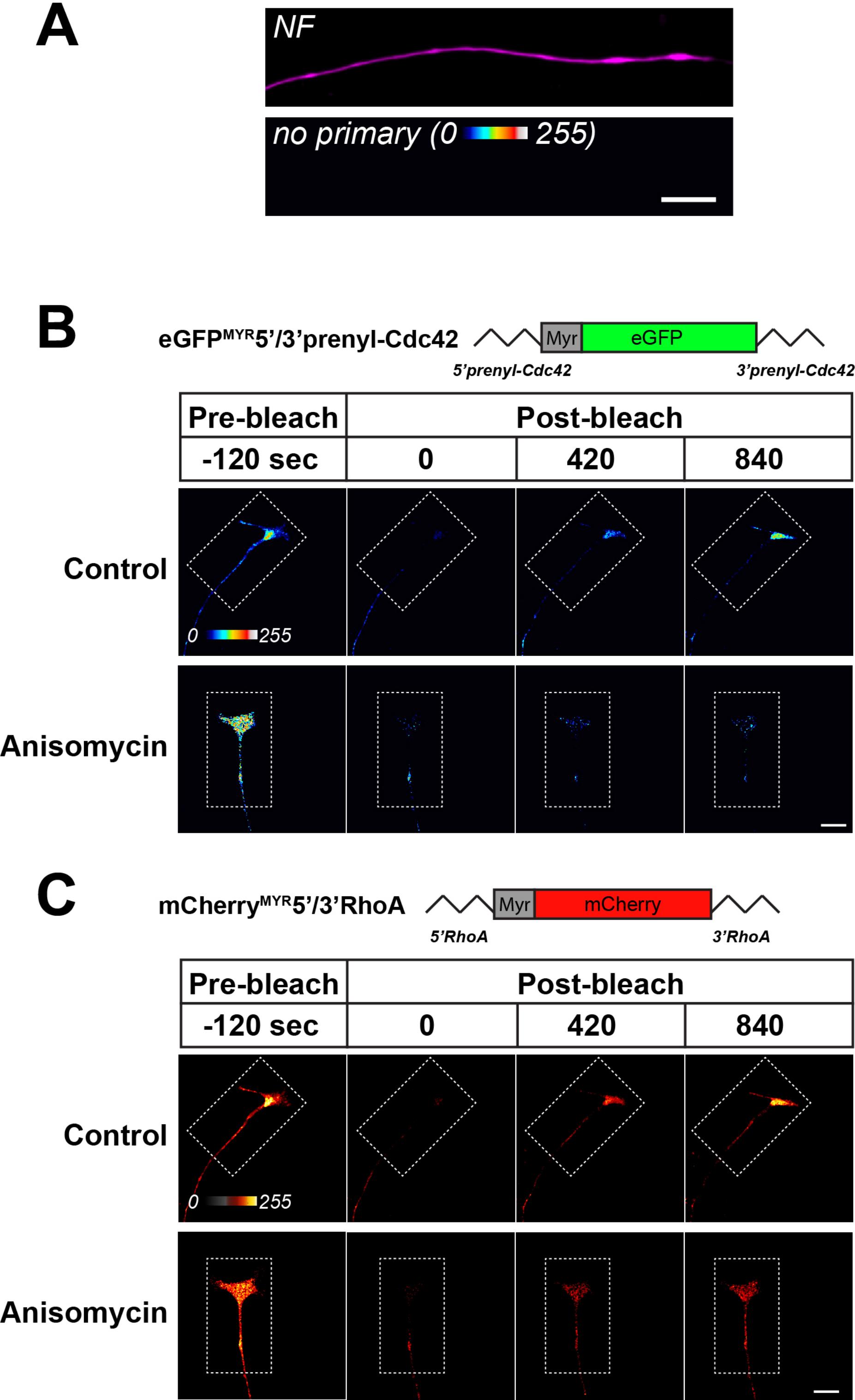

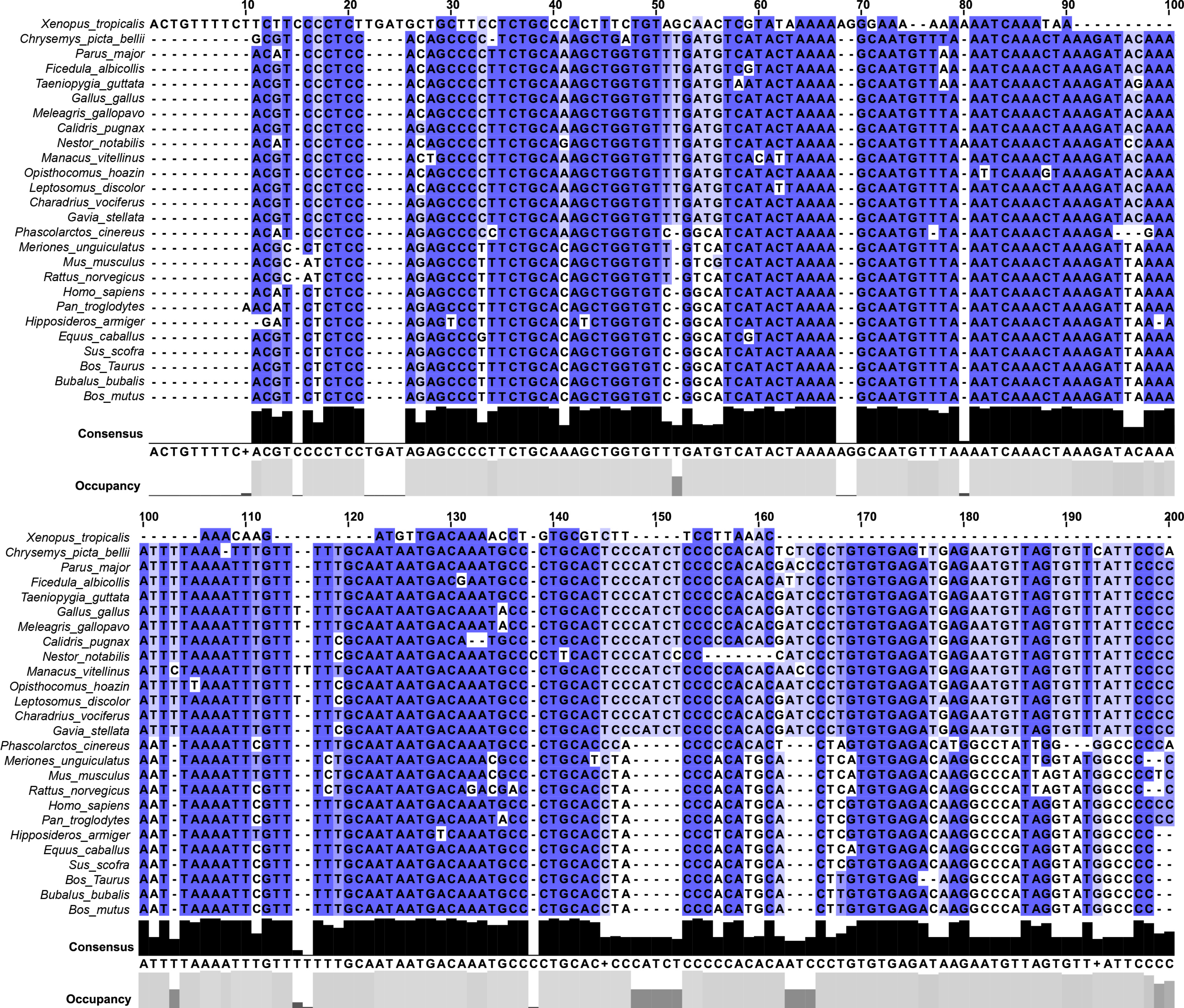

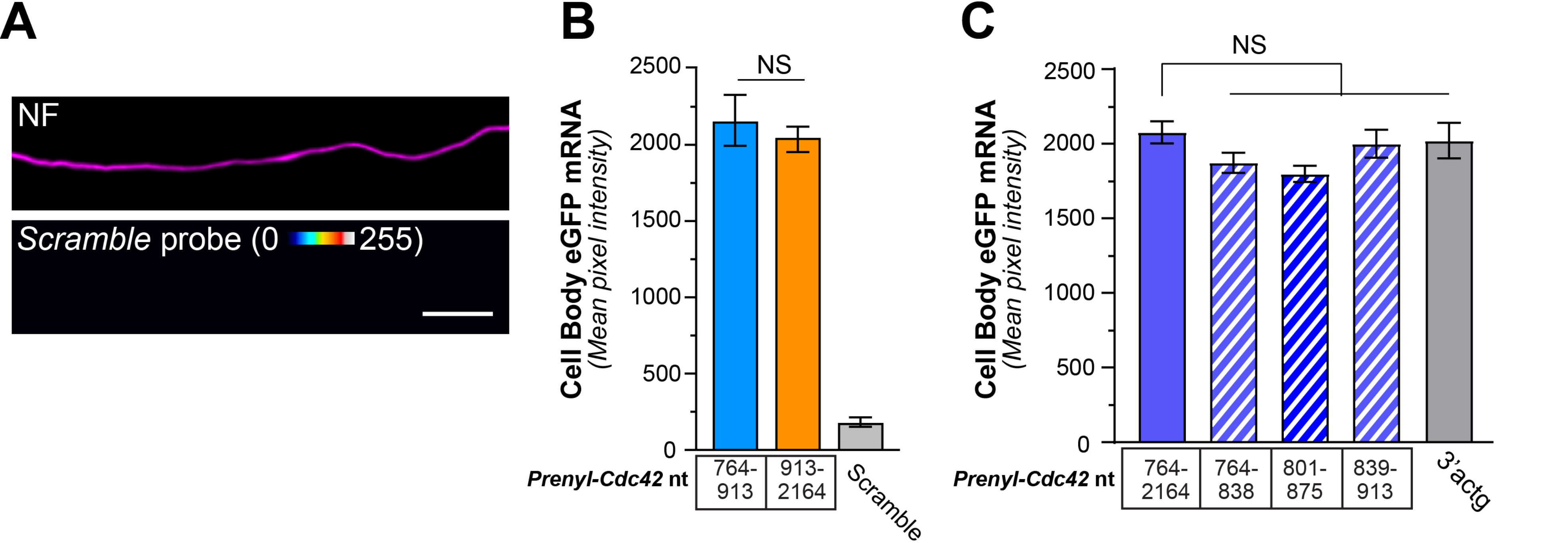

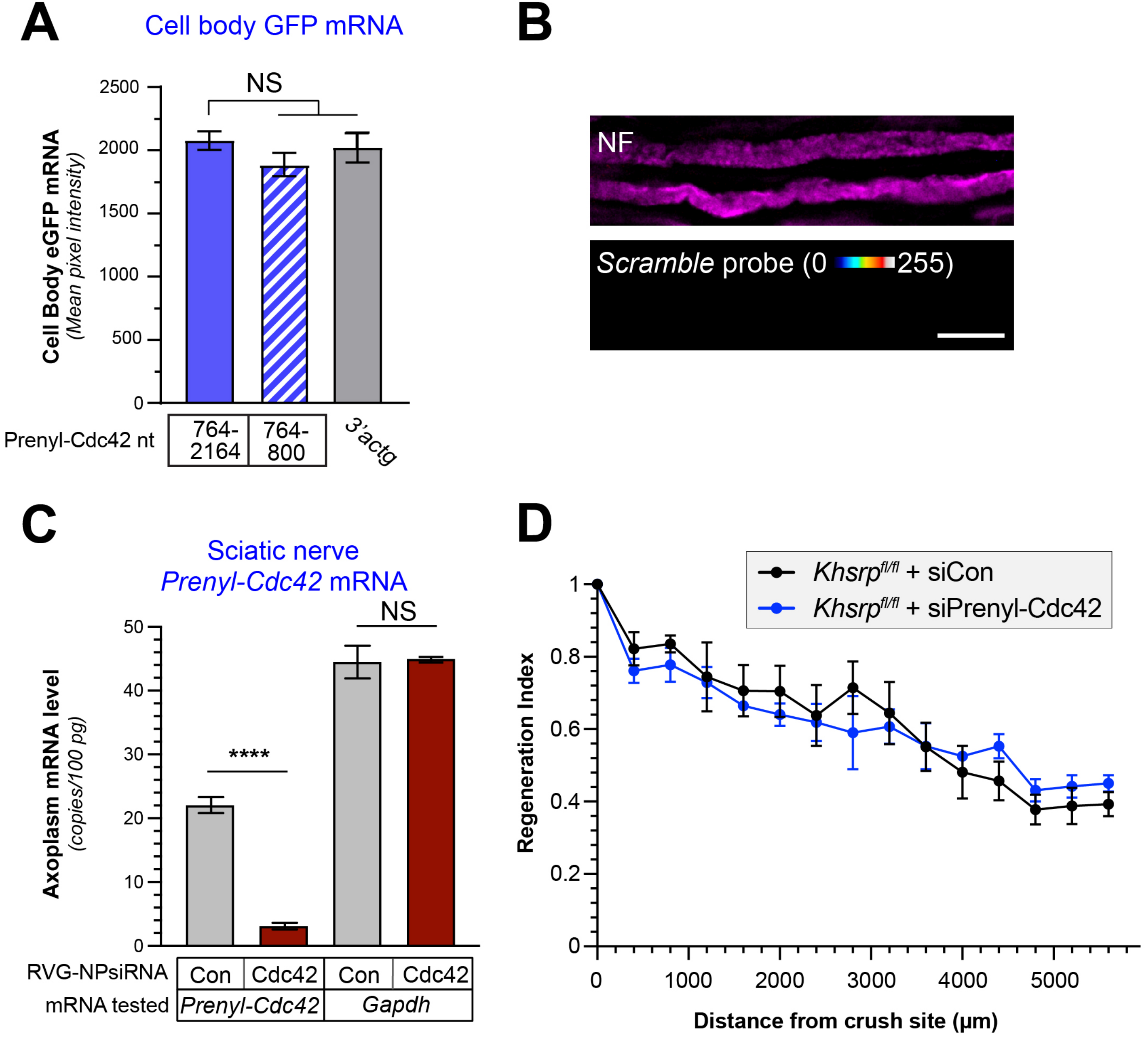

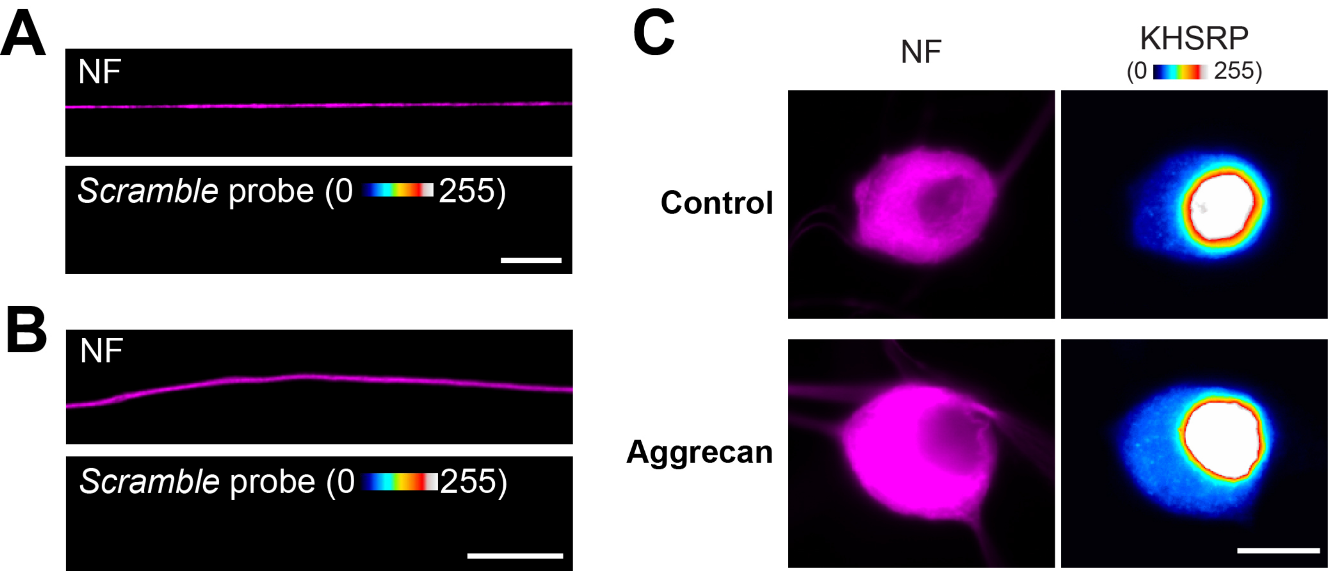
